# Minimal frustration underlies the usefulness of incomplete and inexact regulatory network models in biology

**DOI:** 10.1101/2022.06.07.495167

**Authors:** Shubham Tripathi, David A. Kessler, Herbert Levine

## Abstract

Regulatory networks as large and complex as those implicated in cell-fate choice are expected to exhibit intricate, very high-dimensional dynamics. Cell-fate choice, however, is a macroscopically simple process. Additionally, regulatory network models are almost always incomplete and / or inexact, and do not incorporate all the regulators and interactions that may be involved in cellfate regulation. In spite of these issues, regulatory network models have proven to be incredibly effective tools for understanding cell-fate choice across contexts and for making useful predictions. Here, we show that minimal frustration—a feature of biological networks across contexts but not of random networks—can compel simple, low-dimensional steady-state behavior even in large and complex networks. Moreover, the steady-state behavior of minimally frustrated networks can be recapitulated by simpler networks such as those lacking many of the nodes and edges, and those that treat multiple regulators as one. The present study provides a theoretical explanation for the success of network models in biology and for the challenges in network inference.

## I. INTRODUCTION

Biological network models that describe the regulatory relationship between different molecular players or between higher-level biological entities (such as signaling pathways or cell types) have been extremely useful in systems biology [1, 2] to model and understand the features of cell-fate regulation [3]. With the advent of high-throughput molecular profiling techniques, networkbased models and approaches have become nearly indispensable [4, 5]. Identifying features that distinguish biological networks from random networks has been an area of active research. Previous studies have argued that biological networks present a scale-free degree distribution [6, 7], are hierarchically organized [8], and exhibit recurrence of certain patterns called motifs with a higher probability than expected by random chance [9]. However, these and other analyses of topological differences have provided little insight into the functional differences between actual biological networks and random networks, differences that enable biological networks to effectively regulate cell fates.

Two functional behaviors of biological regulatory networks stand out. First, physics would suggest that even systems with a relatively small number of independent variables are expected to exhibit exceedingly complex behaviors [10, 11]. However, cell-fate regulation, successfully modeled using large and complex networks, is a macroscopically simple process [12–18]. Different cell fates are characterized by distinct expression patterns or activity levels of sets of genes (including transcription factors, micro-RNAs, etc.) [19]. The typical approach to model the establishment of distinct cell fates is to simulate the dynamics of a regulatory network using a methodology of choice (ordinary differential equation-based modeling or rule-based modeling, among others), identify the steady states of network dynamics, and then map each steady state or each group of similar steady states to a distinct cell fate. While the set of cell types—specific gene expression patterns seen in biology—is fairly limited, dynamical models of the size and complexity of biological regulatory networks should, in general, be capable of exhibiting a far more diverse set of expression patterns at steady state. Is there then a universal feature of regulatory networks in biology that restricts the set of gene expression patterns commonly seen?

Second, nearly all network descriptions of cell-fate regulation involve models that are inexact and / or incomplete—such network models do not incorporate all of the genes involved or all the interactions between the chosen genes, and often treat multiple biomolecules as a single regulator. This is a consequence of the limited resolution of current experimental techniques, limited data availability, noise in the collection and interpretation of data from high-throughput experiments, the high context-dependence of biological assays, and, in many cases, choices made to simplify the modeling task. For example, gene networks of widely different sizes have been used to usefully model the regulation of choice between epithelial and mesenchymal cell-fates [13, 14, 20–22]. While none of these network models can claim to be more exact than the others, all can claim to recapitulate the gene expression patterns associated with epithelial and mesenchymal cell fates, and to provide useful insights into the regulation of the underlying biological process. The success of these incomplete and inexact regulatory network models raises the following question: is there a universal feature of regulatory networks in biology that allows us to re-capitulate the observed biological behavior and make useful predictions without the need to know and incorporate the exact network structure?

Our previous work [23] answered, in part, the first question. We identified minimal frustration as a key property of biological regulatory networks across contexts and showed, within a Boolean modeling framework [14], that biological networks exhibit certain steady states with exceptionally low frustration. These states are the ones that are most frequently encountered when simulating network behavior and correspond to the gene expression patterns seen in biology. Such low-frustration states are not seen in the case of random networks that have the same topological features as the biological network. While minimally frustrated biological networks can still exhibit steady states with non-biological gene expression patterns, such steady states are rarely dynamically encountered.

In the present study, we extend our analysis to ordinary differential equation-based models of biological regulatory networks. We show that provided the network is minimally frustrated as defined previously [23], the steady-state network behavior is simple and largely one-dimensional, in spite of the complex and multi-dimensional nature of the network model. This property underlies the suitability of large biological networks for describing a macroscopically one-dimensional process such as cell-fate regulation. We then go on to answer the second question posed above and show that the behavior modeled by a minimally frustrated network can be recapitulated by much smaller, simpler network models either lacking many of the regulators and interactions present in the original network, or combining multiple regulators into single nodes. Thus, the present study builds upon the analysis in [23] to establish minimal frustration as a key feature of biological regulatory networks and helps explain the success of necessarily incomplete systems biology models in modeling cell-fate regulation.

## II. MODELING REGULATORY DYNAMICS

### Specifying regulatory networks

A regulatory network involved in cell-fate regulation can be specified using a directed graph. A node in such a graph may correspond to a transcription factor, a micro-RNA, an epigenetic modifier, or any other regulatory factor. Each directed edge in the graph is signed—either activating or inhibiting—depending on the type of the regulatory relationship between the regulators. Mathematically, a regulatory network with *N* nodes can be described with an *N* × *N* connection matrix *J* such that *J_ij_* = +1 if the edge *i* ← *j* is activating and *J_ij_* = –1 if the edge is inhibitory; *J_ij_* = 0 if there is no edge from *j* to *i*. Provided that the rules governing how the different inputs to a node combine are available, the network dynamics may be simulated either within a discrete modeling framework (a Boolean framework being the most commonly used [24]) or a continuous framework involving ordinary differential equations (ODEs).

### Boolean modeling and definition of frustration

In a Boolean modeling framework, the state of an N-node network is specified by a sequence of *N* binary variables {*s*_1_, *s*_2_,..., *s_N_*}, with *s_i_* = +1 if the regulatory species represented by node i is active and / or highly expressed, and *s_i_* = –1 otherwise. The various inputs to a given node, as described by the connection matrix *J*, may combine additively or via more complex logic-based rules. Here, we consider the simple case wherein the inputs to any given node combine additively and independently [14]. In such a scenario, the discrete-time network dynamics are given as [23]

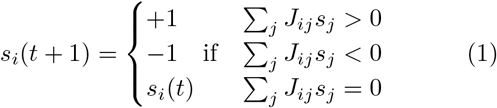

The network state is updated in an asynchronous fashion: at any given point in time, a node is chosen at random and its state updated using Eq. 1. Note that simulating the dynamics of biological regulation using Eq. 1 requires only the network connection matrix *J*—there are no other parameters involved. One can identify the stable states of such dynamics as any network state {*s_i_*} wherein *s_i_*(*t* + 1) = *s_i_*(*t*) for all *i*. For every network state, one can define frustration as the fraction of network edges that are not satisfied in that state, *i.e.*, the fraction of edges for which *J_ij_s_i_s_j_* < 0 [23, 25]. In the case wherein the inputs to a given node combine via logic-based rules, frustration of a network state may be similarly defined. However, the precise mathematical definition is more complex in such a scenario (see [23]).

Throughout this manuscript, we refer to a network as being minimally frustrated if, within the Boolean modeling framework described above, the network exhibits certain steady states with frustration significantly lower than that of the steady states exhibited by random networks with similar topological features (*i.e.*, random networks with the same number of nodes and edges, the same number of activating and inhibitory edges, and the same in-degree and out-degree for each node). Such random networks can be generated from the original biological network by repeatedly choosing a pair of network edges at random and swapping their targets (Appendix Sec. 1). The random networks thus obtained may be more or less modular (as quantified by the directed Louvain modularity [26]) than the corresponding biological network (Fig. S1).

### ODE-based modeling

In an ODE-based modeling framework, the regulatory network state is described by a continuous *N*-dimensional vector {*y*_1_, *y*_2_,..., *y_N_*} where *y_i_* describes the expression or activity level of the regulator represented by node *i*. Given a connection matrix *J*, the network dynamics (in continuous time) can be described using a set of ordinary differential equations of the form [27]

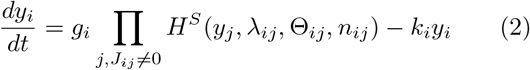

Here, *H^S^* is the shifted Hill function: 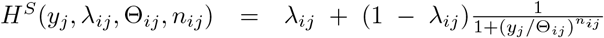. Note that *λ_ij_* > 1 if the edge *i* ← *j* is activating (*i.e.,* if *J_ij_* = +1) and *λ_ij_* < 1 if the edge *i* ← *j* is inhibitory (i.e., if *J_ij_* = –1). Once again, we assume that the inputs to any given node act independently of one another. The system of ODEs in Eq. 2 associates two kinetic parameters with each network node: *g_i_*, the production rate, and *k_i_*, the degradation rate of the regulator represented by node *i*. Three kinetic parameters are associated with each network edge: *λ_ij_*, the maximum fold change in the production rate of node *i* that node *j* can cause, *θ_ij_*, the threshold parameter of the Hill function, and *n_ij_*, the Hill coefficient. The system of ODEs in Eq. 2 describes a general setup to model the dynamics of a system of regulatory nodes that can activate or inhibit one another. Such a system can be defined for any given connection matrix *J*. A more specialized setup with different equations explicitly modeling different modes of biological regulation (*e.g.,* transcriptional regulation, micro-RNA-mediated regulation, ubiquitination-mediated regulation, etc.) may be chosen to model specific biological systems of interest.

For a given network connection matrix *J*, the dynamics in an ODE-based framework will, of course, depend on the choice of the kinetic parameters involved in Eq. 2. The choice of an appropriate parameter set will vary with the biological context and, in general, can be exceedingly difficult. Here, we analyze generic, statistical features of the dynamics for a fixed connection matrix *J* and an ensemble of kinetic parameter sets generated using the random circuit perturbation (RACIPE) approach [27] (see Appendix Sec. 2 for details). RACIPE generates an ensemble of kinetic parameter sets in a systematic fashion such that the ensemble is representative of all biologically relevant possibilities. This approach ensures that our analysis is not restricted to the dynamical behavior under a fixed parameter set fitted to some given (arbitrarily chosen) experimental context. More importantly, it allows us to capture the heterogeneity in dynamical behavior that is inherent in biological systems.

## III. RESULTS

### A. Steady-state dynamics of biological regulatory networks are simple

We analyzed features of the set of steady states exhibited by multiple biological regulatory networks taken from the literature [15, 16, 22] for an ensemble of kinetic parameter sets (as described in Appendix Sec. 2), and compared these features with those obtained for random networks with similar topological features. Fig. 1 a-c show that in the case of biological networks, most of the variation in the steady states is one-dimensional—along the first principal component. This suggests that while these networks are complex—they involve many biomolecular regulators and numerous interactions among them—their behavior at steady state is simple and can be sufficiently described by a single order parameter (*e.g.,* the first principal component). There is no need to specify the expression / activity levels of all of the network nodes to describe the network steady state. In contrast, in the case of random networks, the behavior at steady state is much more complex: a much smaller fraction of the total variance in the set of steady states is captured by the first principal component than in the case of the biological network sharing similar topological features. Thus, describing the steady state in the case of random networks would require specifying the expression / activity levels of many or all of the network nodes, and restricting the description to the first (or even the first few) principal component(s) would be uninformative (Fig. S2). Note that RACIPE was run *ab intio* for each biological network and each random network instance. Thus, the ensemble of kinetic parameters for which the network behavior is simulated is distinct in each case.

**FIG. 1.**
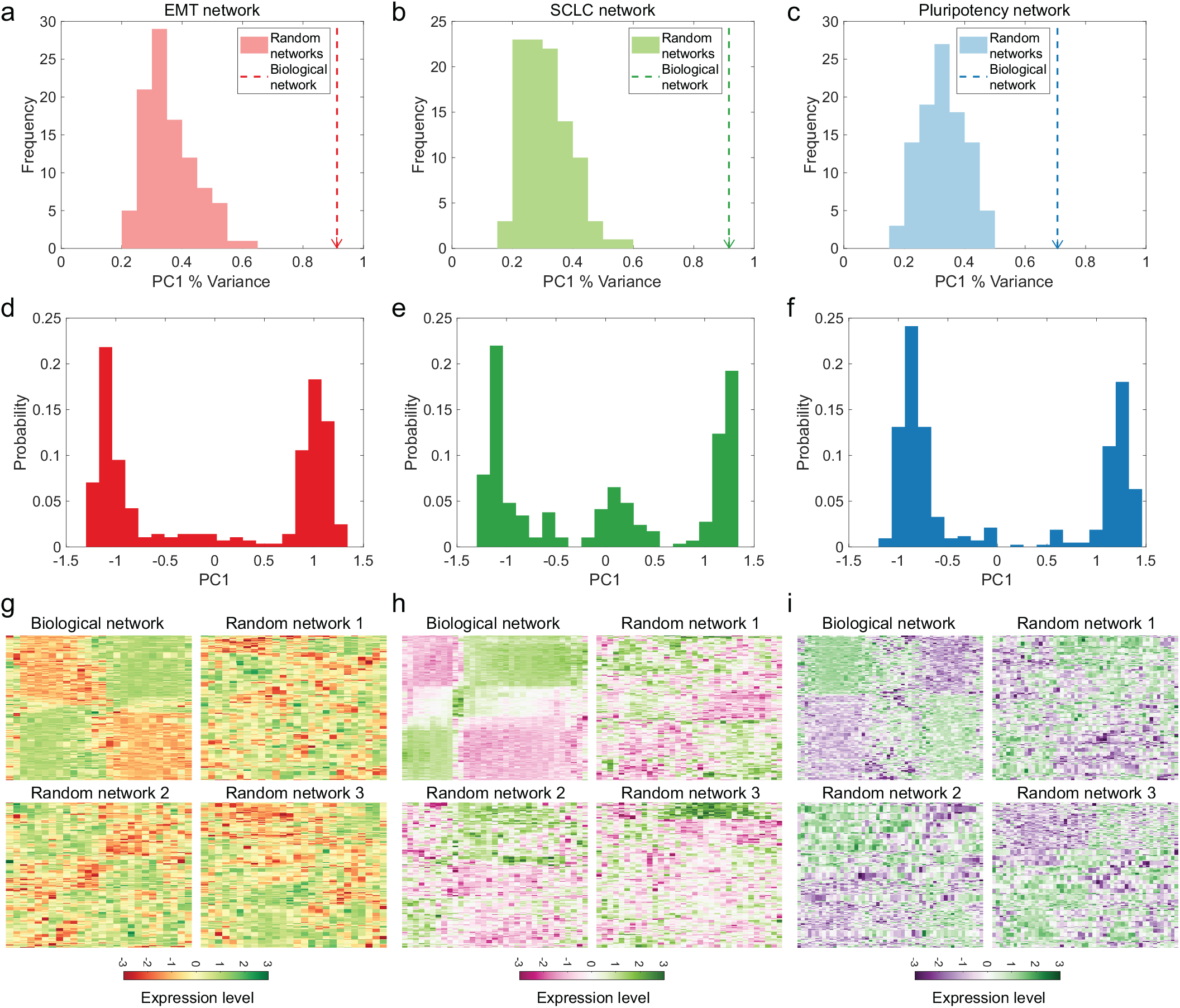
Steady-state dynamics of complex biological networks are simple and largely one-dimensional. **a-c** Distribution of the percentage variance explained by the first principal component (PC1 % variance) in the case of random networks (histograms) with the same number of nodes and edges, and similar topological features as the biological network (dashed vertical lines with arrows). Note that the principal components were sorted in decreasing order of the percentage variance explained. Thus, in each case, the first principal component (PC1) is the one that explains the greatest percentage of the variance in the steady states. The sets of steady states for biological and random networks were obtained using the approach described in Appendix Sec. 2. See Fig. S2 for the fraction of variance in the steady states explained by the other principal components. **d-f** Distribution of the first principal component (PC1) projection of the steady states obtained for the three biological networks. See Fig. S3 for the projection of the steady states along the first and second principal components. **g-i** Expression levels of the different network nodes in the various steady states obtained for biological and random networks. Different steady states are shown along the rows while the network nodes are shown along the columns of the heatmap, with the colors indicating the expression levels. Both rows and columns were hierarchically clustered to obtain the heatmap in each case (see Appendix Sec. 3). The left column (panels **a**, **d**, and **g**) shows the behavior in the case of the epithelial-mesenchymal transition (EMT) network [22], the middle column (panels **b**, **e**, and **h**) shows the behavior in the case of the SCLC network [16], and the right column (panels **c**, **f**, and **i**) shows the behavior in the case of the pluripotency network [15].

Analyzing the underlying structure in the set of steady states obtained in different cases, we find that the distribution of the first principal component in the case of the biological networks analyzed here is largely bimodal (Fig. 1 d-f). This indicates that the steady states obtained for the ensemble of parameter sets in the biological case cluster into two distinct groups. This observation is confirmed visually by hierarchical clustering of the set of steady states obtained (top-left plot in Fig. 1 g-i) wherein we see two sets of steady states with distinct activity patterns of the network nodes (also, see Fig. S3). No such discernible pattern can be seen in the case of random networks (plots other than the top-left one in Fig. 1 g-i). Note that each parameter set in the ensemble generated by RACIPE may be interpreted as modeling a different cell in a population [27], with the variation in the ensemble capturing the cell-to-cell variation in the population. The readily evident clustering of the steady states into two distinct groups (Fig. 1 d-i) indicates that most of the steady states of these networks can be mapped to one of two phenotypic states with distinct gene expression patterns. This is consistent with the role of these networks in establishing two distinct cell fates: epithelial cells and mesenchymal cells in the case of the epithelial-mesenchymal transition (EMT) network [22], neuroendocrine cells and mesenchymal cells in the case of the small cell lung cancer (SCLC) network [16], and stem cells and differentiated cells in the case of the pluripotency network [15].

Recall that the only input to RACIPE is the connection matrix *J*. The approach does not take any other experimental data as input, generating the ensemble of kinetic parameters in an unbiased fashion so as to capture the range of possible network behaviors. Thus, the structure in the steady states obtained using RACIPE (shown in Fig. 1 d-i) is an intrinsic property of biological connection matrices that is seen to be absent in random networks.

Fig. 1 demonstrates the capability of the biological networks analyzed here to robustly establish cell types with biological gene expression patterns. Biological networks exhibit this behavior for a broad range of kinetic parameter sets in a manner that is dependent on the network connection matrix *J*. While these networks can still exhibit certain steady states that cannot be uncontroversially mapped to one of the two groups that correspond to biological phenotypic states (Fig. 1 d-f) and with expression patterns not seen in canonical cell types, such steady states are infrequently encountered when simulating network dynamics starting from random initial conditions. Such steady states with aberrant expression patterns are seen at a higher frequency in the case of the SCLC network (see Fig. 1 e, h) as has been noted elsewhere [16]. The non-canonical expression patterns corresponding to such aberrant steady states, while suppressed in healthy cells, have been reported in cancer cells [16, 28].

### B. Minimal frustration underlies the simple steady-state dynamics of biological regulatory networks

We have previously demonstrated that biological regulatory networks taken from the literature (including the ones analyzed in Fig. 1), within a Boolean modeling framework, exhibit certain steady states with frustration lower than that of steady states exhibited by random networks with similar topological features, *i.e.*, biological regulatory networks are minimally frustrated [23]. In the previous section (Sec. III A), we have shown that, within an ODE-based modeling framework, the steady states exhibited by biological networks are simple and largely one-dimensional. In both cases, we argue that the reported biological network behavior underlies the ability of large and complex networks to describe cell-fate regulation. To determine if the two features—minimal frustration within a Boolean modeling framework and simple, largely one-dimensional steady-state dynamics within an ODE-based modeling framework—are directly related, we simulated the evolution of a population of random networks with the same topological features as the EMT network [22] subject to different selection pressures (see Appendix Sec. 4 for the detailed methodology). Under selection for networks for which the steady-state dynamics are largely one-dimensional (as quantified by the percentage of variance in the set of steady states explained by the first principal component) (Fig. 2 a), we obtained networks that were minimally frustrated (Fig. 2 b). Reciprocally, selection for the low-frustration property (Fig. 2 c) led to the emergence of networks with largely one-dimensional steady-state dynamics (Fig. 2 d). Moreover, under selection for low frustration, we obtained networks that exhibited steady-state gene expression patterns very similar to the biological case (compare Fig. 2 f and the top-left plot in Fig. 1 g). It has previously been suggested that a large, complex regulatory network can exhibit lowdimensional gene expression patterns if the network has a modular topology [29]. However, we did not see an increase in the modularity of networks in the population while selecting for networks with largely one-dimensional steady-state dynamics (Fig. S4).

**FIG. 2.**
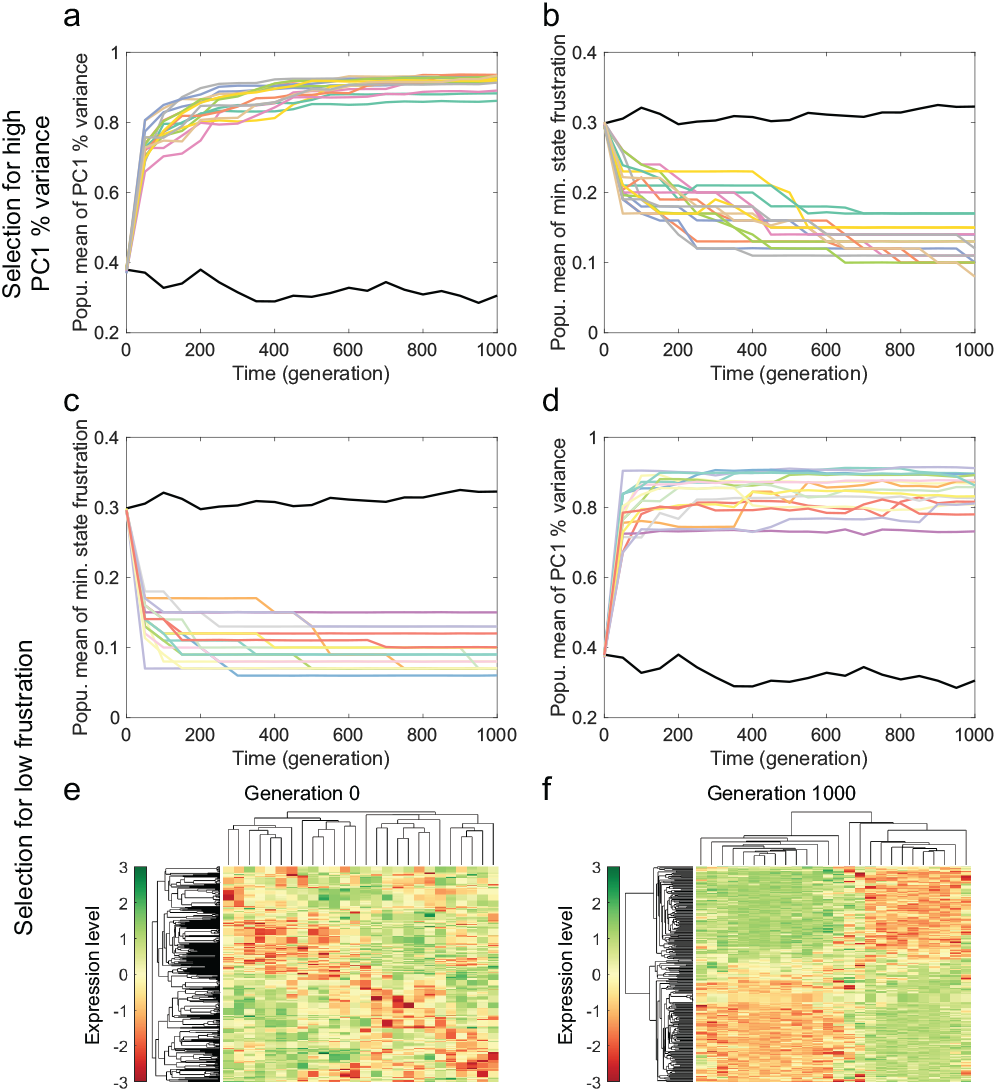
Selection for networks with simple, one-dimensional steady-state dynamics automatically selects for minimal frustration, and vice versa. **a-b** Results of an evolution simulation wherein networks for which a larger fraction of the variance in steady-state behavior can be explained by a single principal component (*i.e.*, networks that exhibit simpler, more onedimensional steady-state dynamics) are selected for at each generation. **c-f** Results of an evolution simulation wherein networks with minimally frustrated steady states are selected for at each generation. Panel **e** shows the expression levels of network nodes at steady state for a network randomly chosen from the population at generation 0. Panel **f** shows the expression levels at steady state for a network randomly chosen from the population at generation 1000. In panels **a** and **c**, the mean population PC1 score is shown. In panels **b** and **d**, the mean population frustration score is shown. Both these scores are defined in Appendix Sec. 4. In panels **e**-**f**, different steady states (obtained using RACIPE; see Appendix Sec. 2) are shown along the rows while the network nodes are shown along the columns of the heatmap. Expression levels are indicated by the color (see adjacent color bars). Both rows and columns were hierarchically clustered to obtain the heatmaps. In panels **a**-**d**, the black curve shows the behavior for an evolution simulation in the absence of any selection pressure. The other colors indicate independent simulation runs (with selection). See Appendix Sec. 4 for details of the simulation setup.

The behavior in Fig. 2 shows that the minimal frustration property and the property of exhibiting simple, largely one-dimensional steady-state dynamics over an ensemble of parameter sets are, in fact, equivalent—selection for one automatically selects for the other. Since a Boolean model, defined here simply by the connection matrix *J,* can be built into a corresponding ODE-based model by including a suitable set of kinetic parameters and mathematical expressions, we may conclude that the minimal frustration property within the Boolean framework underlies the simple steady-state dynamics seen in the ODE-based framework. An approach for directly obtaining a Boolean modeling framework starting from an ODE-based model would be helpful for verifying if simple steady-state dynamics in an ODE-based model can underlie minimal frustration within a Boolean modeling framework. Such an approach will be investigated in a future study.

### C. Simplicity of steady-state dynamics is preserved under node and edge deletions

Robustness of functional behavior to genomic and environmental perturbations is a well-known feature of biological systems [30]. To determine if the functional characteristic of biological regulatory networks highlighted here—simple, largely one-dimensional steadystate dynamics—is robust to node and edge deletion, we deleted nodes (Fig. 3 a) and edges (Fig. 3 b) in the EMT network [22] one-by-one (following the approach detailed in Appendix Sec. 5), and reported the percentage of variance in the steady states that is explained by the first principal component (corrected for the number of nodes in the network) at each step. Fig. 3 a-b show that the steady-state dynamics remain largely onedimensional even as nodes and edges are successively deleted from the EMT network, and this behavior is maintained over the deletion of a large fraction of nodes and edges. Similar behavior is observed in the case of a minimally frustrated network (Fig. 3 c-d) obtained at the end of the evolution simulation subject to selection for low frustration shown in Fig. 2 c, as well as for other biological networks (Fig. S5). The shape of the distribu-tion of the first principal component can also withstand the deletion of a large fraction of network nodes and edges (Fig. 3 e-f and Fig. S6). Note that the change in the variance along the first principal component depends on the order in which the nodes or edges are deleted (compare the red (blue) plots with the pink (light blue) plots in Fig. 3 a-d), indicating that certain nodes and edges in the network are more important that others in maintaining the simple, one-dimensional network dynamics.

**FIG. 3.**
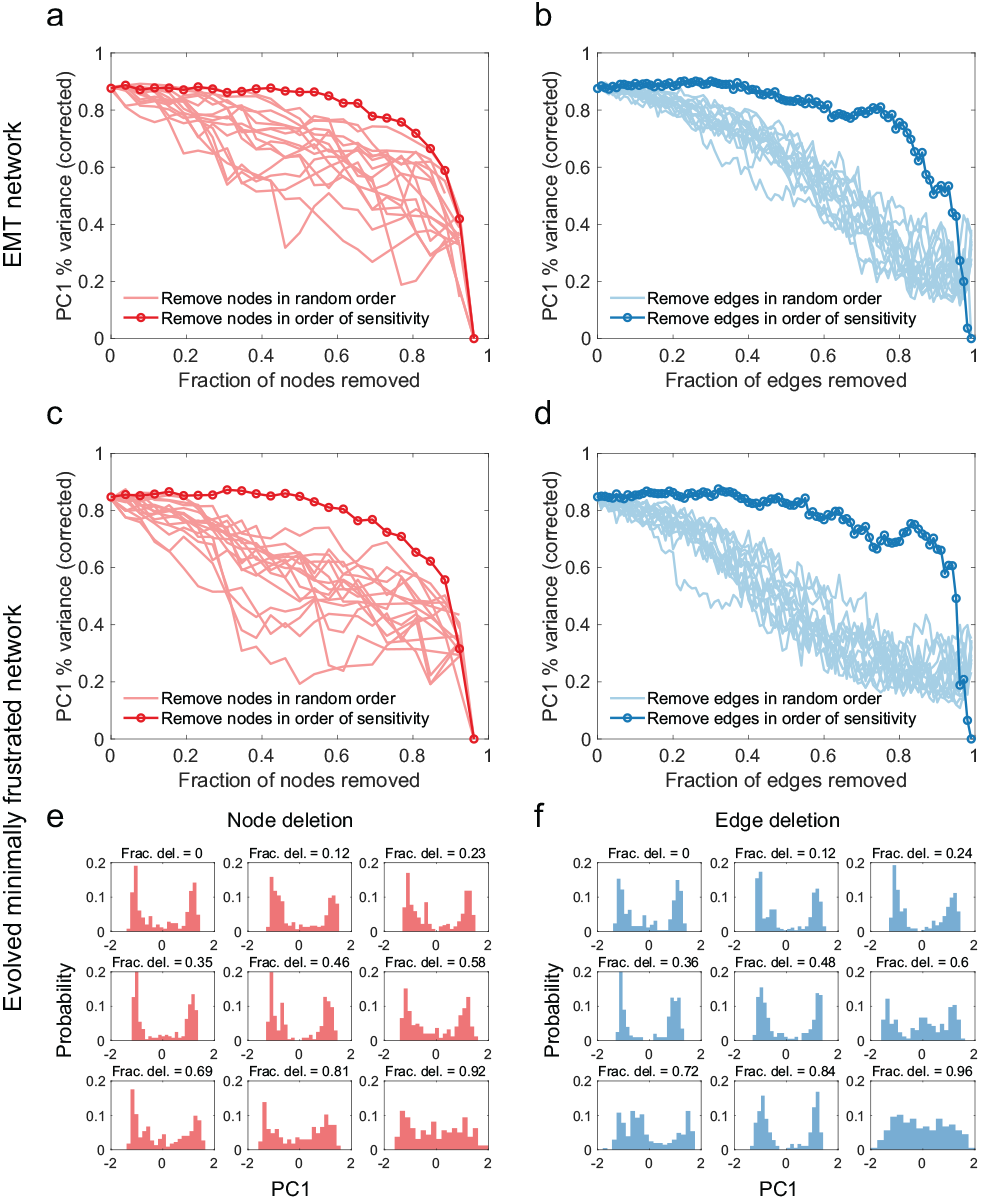
The simple, one-dimensional steady-state behavior (as characterized by a large fraction of the variance in the steady states being explained by the first principal component (PC1)) is preserved under node and edge deletion in the case of the 26 node, 100 edge epithelial-mesenchymal (EMT) network [22] (**a-b**) as well as in the case of a minimally frustrated network obtained via an evolution simulation (**c-f**). **e-f** Change in the distribution of the first principal component (PC1) projection of the steady states during node (**e**) and edge (**f**) deletion. We see that a large fraction of the nodes (**e**) and edges (**f**) in the original network can be deleted before the distribution of the first principal component deviates significantly from the distribution in the case of the original network (shown in the top-left plots in **e** and **f**). See Fig. S6 for a quantitative comparison of the distributions obtained after node or edge deletion with the original distribution.

### D. Networks lacking multiple nodes and edges can recapitulate biological expression patterns

Until now, we have shown that in spite of their large size and complexity, minimally frustrated networks such as biological networks taken from the literature exhibit fairly simple, one-dimensional steady-state dynamics. Based on this observation, we hypothesized that “simpler” network models should be capable of recapitulating the steady-state behavior exhibited by a larger, more complex network provided the larger network is minimally frustrated. Here, by “simpler” we imply networks lacking multiple nodes and / or edges present in the original network. A different manner of network simplification is addressed in the next section.

In agreement with the above hypothesis, we find that networks lacking numerous nodes and edges present in the EMT network [22] (see Appendix Sec. 5 for how such networks were obtained) can still recapitulate the pattern of node expression / activity levels exhibited by the full 26 node, 100 edge EMT network (Fig. 4 a, c). Importantly, these smaller, simpler networks exhibit strikingly similar behavior in response to gene knockouts as the full EMT network (Fig. 4 b, d): the change in the distribution of the first principal component upon gene knockout in the simpler networks is qualitatively similar to the change in the case of the original network. Thus, given experimental data on the gene expression profiles seen in cells and even data on the effect of knocking out multiple genes, it is impossible to identify any one network model as uniquely the correct one. Instead, one can employ many useful network models of different sizes and varying complexities to model EMT, each missing a different subset of the nodes and edges present in the original network. Clearly, it is not necessary to know the exact network to recapitulate the overall biological behavior. In fact, the 26 node, 100 edge EMT network considered as the full EMT network in this section is itself not the “correct” network—while it captures many of the features of EMT regulation, it cannot claim to incorporate all the regulators that can affect EMT or even the entire set of interactions among the regulators it does incorporate. Note that useful network models of EMT cannot be simplified beyond a certain limit: exceedingly simple networks cannot adequately recapitulate biological behavior. This can be clearly seen in Fig. 4 c. Interestingly, the behavior described above is not limited to the EMT network. Fig. S11 shows that for a minimally frustrated network obtained via the evolution simulation (Fig. 2 cd), the steady-state node activity pattern can also be recapitulated by simpler networks with fewer nodes and edges.

**FIG. 4.**
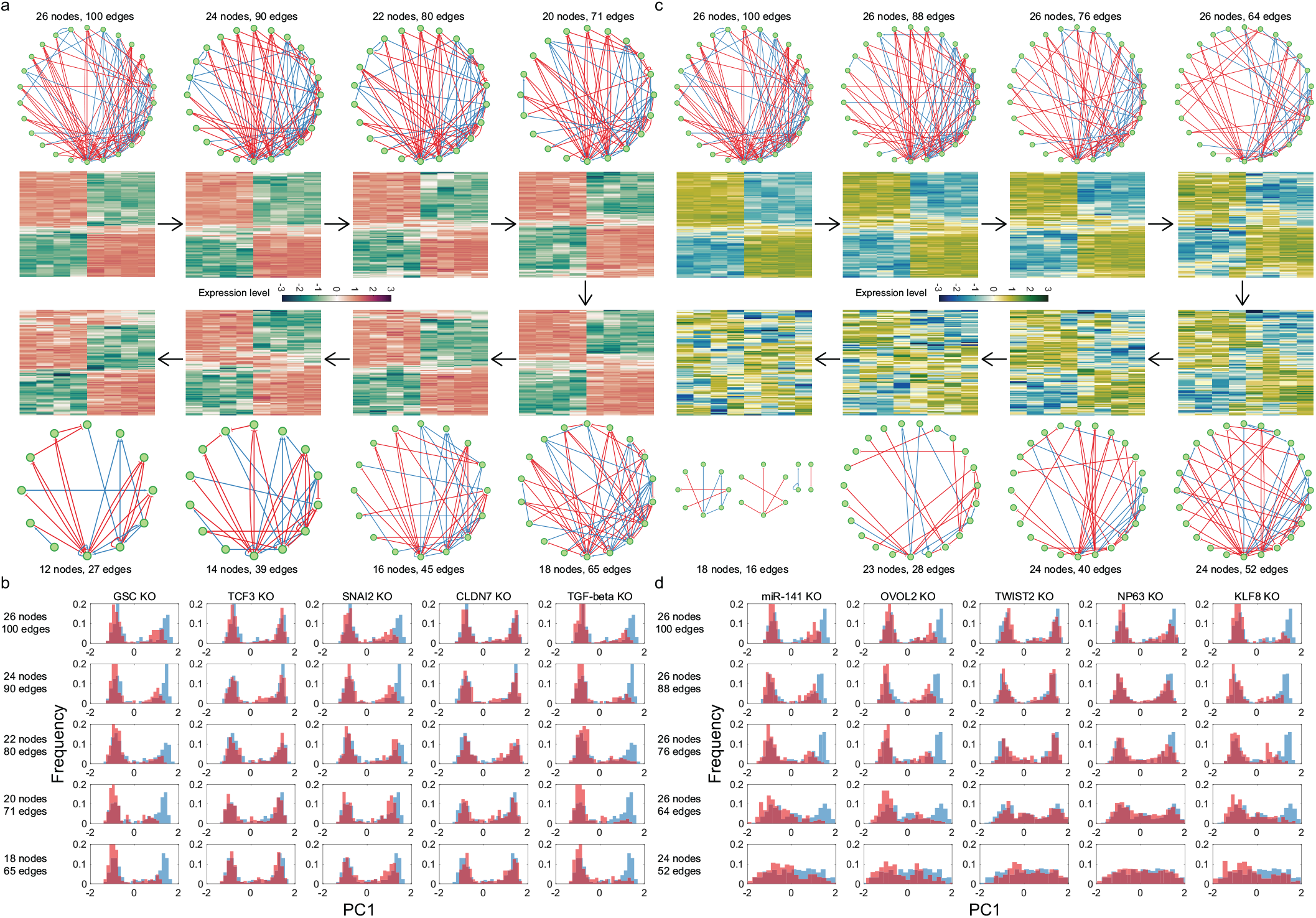
The pattern of node expression / activity levels at steady state exhibited by the 26 node, 100 edge epithelial-mesenchymal (EMT) network [22] is recapitulated by simpler networks obtained upon node deletion (panel **a**) or edge deletion (panel **c**). Each heatmap shows the expression levels (indicated by the color) of the same eight nodes of interest across the different steady states obtained for each network. Different steady states are shown along the rows while the network nodes are shown along the columns of the heatmaps. The heatmaps shown here were generated by hierarchically clustering the rows (*i.e.*, the steady states). Each heatmap shows the network nodes (*i.e.*, the columns) in the same order. The simpler networks obtained are shown alongside the corresponding heatmaps. **b, d** The simpler networks obtained upon node or edge deletion recapitulate the response of the larger, original EMT network to multiple gene knockouts. In each plot shown in **b** and **d**, the blue histogram shows the distribution of the first principal component in the control case while the pink histogram shows the distribution obtained upon gene knockout. The principal component analysis was carried out for the eight nodes of interest. Each row in panels **b** and **d** shows the behavior for a fixed network (whose size is indicated in the figure) while each column shows the response to a given gene knockout (KO). See Fig. S7 for a quantitative comparison of the distributions obtained after gene knockout with the distributions in the control case. The simpler networks analyzed here were obtained by successively deleting randomly chosen nodes (panels **a** and **b**) or edges (panels **c** and **d**) while ensuring that the eight nodes of interest are retained in the simpler networks. The same set of eight nodes were the nodes of interest in panels **a**-**d**: four epithelial state markers (CDH1, miR-200b, miR-200c, miR-34a) and four mesenchymal state markers (VIM, ZEB1, SNAI1, and TWIST1).

### E. Networks that combine regulators can recapitulate biological behavior

In the previous section, we analyzed the steady-state behavior of simpler networks that lack many of the nodes and edges present in the larger, original network. Here, we consider simpler networks obtained by combining sets of nodes in the original network and treating each set as a single regulator. Such a network simplification was motivated by the observation that in the literature, cell types and cell-state transitions have been described both in terms of the expression levels of individual genes as well as in terms of the overall activity levels of different pathways (often comprising numerous genes) [31, 32]. We first developed a systematic procedure to combine sets of nodes into a single regulator (described in Appendix Sec. 6). Fig. S8 shows the sequence of networks obtained by repeatedly applying this “coarse-graining” procedure to the EMT network [22]. Note that the network obtained at each step has both fewer nodes and edges, and is thus simpler than the network in the previous step.

Consistent with the behavior in Fig. 4, we find that the simpler networks obtained by applying the abovementioned coarse-graining procedure to the 26 node, 100 edge EMT network are able to recapitulate the steadystate expression patterns exhibited by the original network (Fig. 5 a). Fig. S9 shows two of the simplified networks obtained via the coarse-graining procedure. Note that the network shown on the right therein groups together various epithelial factors (miR-101, miR-200a, miR-141, CLDN7, OVOL2, GRHL2, miR-30c, miR-9) into a single regulatory node and various mesenchymal factors (FOXC2, ZEB2, SNAI2, TGF-beta, TWIST2, GSC, KLF8, TCF3, miR-205, and NP63) into another regulatory node, instead of treating each of these factors separately. This simplified network exhibits a steadystate expression pattern very similar to the one exhibited by the original, larger EMT network (compare the first and last panels in Fig. 5 a). We also applied our coarse-graining procedure to a larger EMT network taken from the literature, one with 72 nodes and 142 edges [14]. Like other biological networks analyzed in the present study, this network is also minimally frustrated [23]. Once again, the simpler networks obtained by coarse graining reproduced the steady-state expression patterns of the key EMT-related genes (Fig. 5 b) exhibited by the larger, original network. The grouping of different nodes in the simpler networks is biologically interpretable (see the network on the right in Fig. S10): genes that are in the same pathway are grouped together. The Hypoxia stimulus, HIF1 gene, and LOXL gene are grouped to form a node representing the hypoxia pathway. Various factors involved in growth-factor signaling, including platelet-derived growth factor (PDGF) and receptor (PDGFR), epithelial growth factor (EGF) and receptor (EGFR), insulin-like growth factor (IGF1) and receptor (IGF1R), and fibroblast growth factor (FGF) and receptor (FGFR), are also grouped to form a single node representing the many signaling pathways known to drive EMT. The Notch pathway genes and factors (NOTCH, NOTCH intracellular domain (NOTCHic), DELTA, Jagged, and HEY1) are grouped together, and so are the molecular players involved in the Wnt signaling pathway (TCF / LEF, Wnt, Frizzled, and AXIN2).

**FIG. 5.**
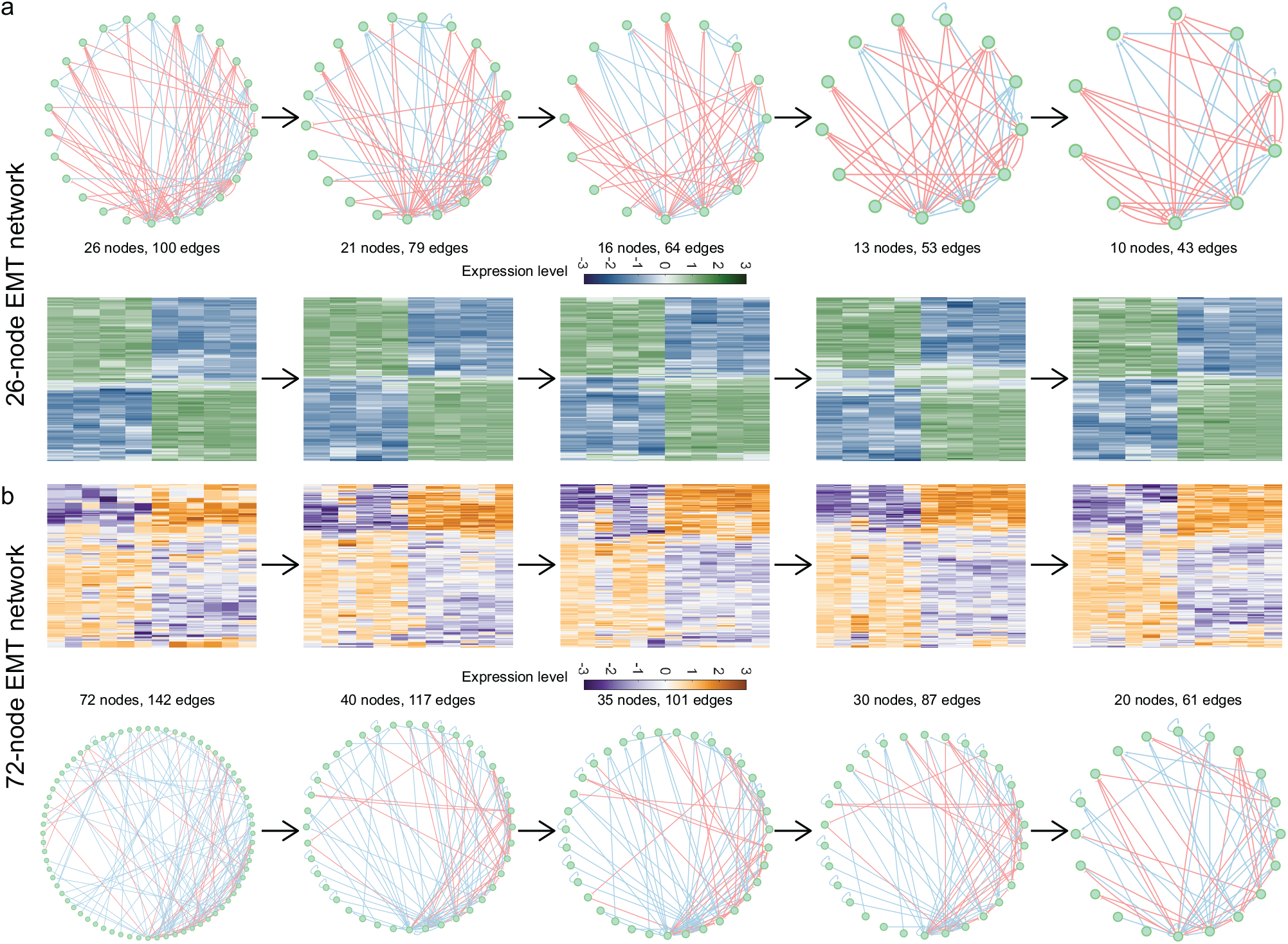
**a** (Bottom) Simpler networks obtained by repeatedly applying the coarse-graining procedure described in Appendix Sec. 6 to the 26 node, 100 edge epithelial-mesenchymal (EMT) network [22] recapitulate the steady-state expression patterns exhibited by the larger, original network. Each heatmap shows the expression levels (indicated by the color) of the same eight nodes of interest (same as the nodes of interest in Fig. 4) across the different steady states obtained for each network. (Top) The original 26 node, 100 edge EMT network is shown (left) along with the simpler networks obtained via the coarse-graining procedure. **b** (Top) Simpler networks obtained via the coarse-graining procedure applied to the 72 node, 142 edge EMT network [14] exhibit steady-state expression patterns similar to the pattern obtained for the larger, original network. Each heatmap shows the expression levels (indicated by the color) of the same twelve nodes of interest across the different steady states obtained for each network. The twelve nodes of interest were Ecadherin, KLF4, cateninmemb, miR200, GSK3, TrCP, cateninnuc, ZEB1, SNAI1, TWIST1, ZEB2, and FOXC2. (Bottom) The original 72 node, 142 edge EMT network is shown (left) along with the simpler networks obtained via the coarse-graining procedure. In both **a** and **b**, the coarse-graining procedure was applied while ensuring that the nodes of interest are not combined with any other network node at any step. In all heatmaps, different steady states are shown along the rows while the network nodes are shown along the columns of the heatmaps. The heatmaps shown here were generated by hierarchically clustering the rows (*i.e.,* the steady states). Each heatmap shows the network nodes (*i.e.*, the columns) in the same order.

Fig. S11 shows that the ability of simpler, coarse-grained networks to recapitulate the steady-state gene expression patterns exhibited by the larger network is not limited to the case of biological networks taken from the literature but extends to minimally frustrated networks obtained via the evolution simulation.

### F. A data-driven example

So far, we have analyzed the behavior of previously published biological networks that were constructed by aggregating information from the literature and from biological databases using a variety of methods [14–16, 22]. For our final example, we turn to a network constructed directly from gene expression data.

#### Regulation of MYC-pathway activation in breast tumors

Terunuma *et al.* obtained the bulk gene expression profiles from 61 breast tumor samples and identified a subset of tumors that showed a MYC-activated phenotypic state [33]. This phenotype was associated with elevated levels of the oncometabolite 2-hydroxyglutarate, DNA hypermethylation, and poor disease prognosis. We used the gene expression data from this study as an input to the GRNBoost2 algorithm [34] and obtained a regulatory network that may be involved in the regulation of tumor cell-fate choice between high MYC-activation and low MYC-activation states (see Appendix Sec. 7 for the detailed methodology). The inferred regulatory network consists of 138 nodes and 451 edges. We have previously shown that such an inferred network is minimally frustrated, just like the biological networks taken from the literature [23]. Here, we report that the inferred regulatory network modeled using ODEs exhibits steady states that vary mostly along the first principal component (Fig. 6 b; red dot), once again consistent with the behavior seen in the case of biological networks taken from the literature (Fig. 1). This was not true for the case of networks inferred by using randomly shuffled gene expression patterns as input to our network inference methodology (Fig. 6 b; blue dots). Importantly, simulation of dynamics of the inferred network using RACIPE (Appendix Sec. 2) recapitulated the gene expression patterns seen in patient tumor samples (Fig. 6 a, d). As in the case of biological networks taken from the literature, the steady-state gene expression patterns exhibited by the large inferred network could be recapitulated by simpler networks obtained by node deletion, edge deletion, or via the coarse-graining procedure (Fig. 6 d). Interestingly, the simpler networks obtained via the coarse-graining procedure better preserve the steady-state expression patterns of the nodes of interest as compared to the simpler networks obtained via node or edge deletion.

**FIG. 6.**
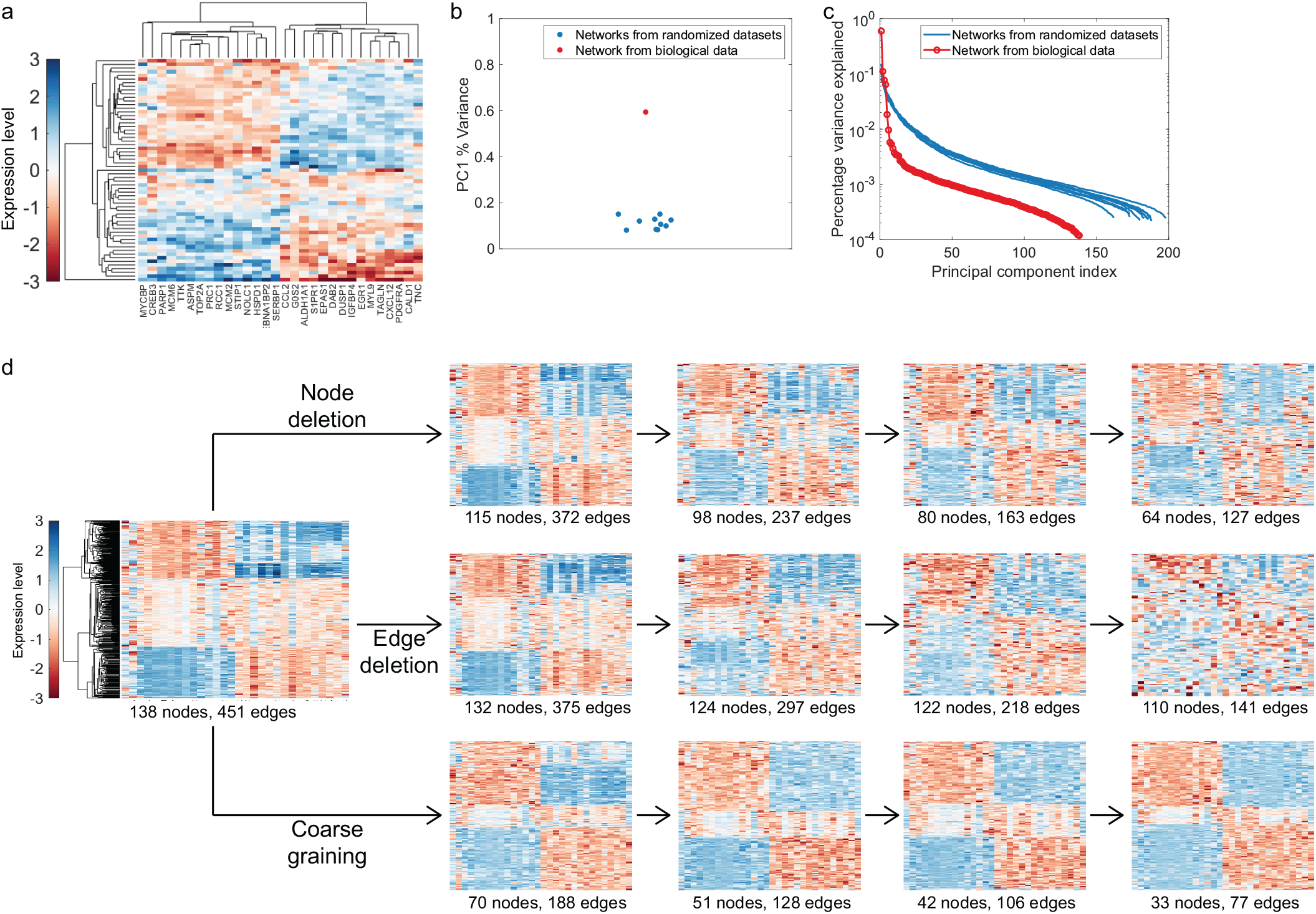
A network inferred from a gene expression dataset [33] exhibits the same behavior as biological networks taken from the literature and other minimally frustrated networks. **a** Expression of 30 MYC-associated genes in 61 breast tumor samples obtained by Terunuma *et al.* [33]. Different tumor samples are shown along the rows while the genes are shown along the columns. Color indicates the expression level. **b** Fraction of variance in the steady states explained by the first principal component (PC1) in the case of the network inferred from the gene expression profiles obtained from breast tumor samples [33] (red dot) and in the case of networks inferred from randomized gene expression profiles (blue dots). Note that the principal components are sorted in decreasing order of the fraction of variance explained. Thus, the first principal component is the one that explains the greatest fraction of the variance in the steady states. **c** The percentage of variance in the steady states explained by the different principal components in the case of the network inferred from a biological dataset [33] (pink curve) and in the case of networks inferred from randomized gene expression datasets (blue curves). **d** (Left) The network inferred from the expression profiles of breast tumor samples [33] exhibits steady states with gene expression patterns similar to those seen in the breast tumor samples in **a**. (Right) Simpler networks obtained via node deletion (top row), edge deletion (middle row), or coarse graining (bottom row) recapitulate the expression patterns exhibited by the larger inferred network. Each heatmap shows the expression levels of the same 30 nodes of interest across the different steady states. The simpler networks were obtained by deleting nodes and edges in random order while retaining the 30 nodes of interest at each stage. While applying the coarse-graining procedure, the 30 nodes of interest were not at any stage combined with any other network node. In all heatmaps, different steady states are shown along the rows while the network nodes are shown along the columns of the heatmaps. The heatmap in **a** was obtained by hierarchically clustering both the rows (*i.e.*, the patient expression profiles) and the columns (*i.e.*, the genes of interest). The heatmaps in **d** were generated by hierarchically clustering the rows (*i.e.*, the steady states): each heatmap shows the network nodes (*i.e.*, the columns) in the same order as in **a**. The names of the 30 nodes of interest are listed below the heatmaps in **a** and **d** (right).

The result shown in Fig. 6 indicates that the behaviors described in the present study do not apply just to chosen biological networks taken from the literature or to idealized minimally frustrated networks obtained via theoretical exercises. The behaviors reported here extend to a biological network inferred from real biological data—here, the gene expression profiles of patient breast tumor samples.

## IV. DISCUSSION

Large, complex networks involving numerous molecular players and various regulatory relationships among them are frequently used to describe and model cell-fate choice in biology, especially as high-throughput assays have become increasingly commonplace. From a dynamical systems perspective, such networks would be expected to exhibit complex behaviors in very highdimensional spaces [10, 11]. However, in most cases, cell-fate choice appears to be a macroscopically simple and low-dimensional process [12], and where a given cell falls on the spectrum between distinct cell fates can be specified with a single order parameter. For example, pseudotime, which is essentially a one-dimensional order parameter, has been widely used to order cells along a cell-fate trajectory based on their gene expression patterns [35, 36]. Multiple studies have described metrics, a single number in each case, to specify where a sample lies on the epithelial-mesenchymal spectrum on the basis of the gene expression in that sample [37–39]. Another one-dimensional metric has been developed to assess the stemness of leukemia samples [40]. In the present study, we have shown that large, complex networks can exhibit largely one-dimensional steady-state behavior provided the network is minimally frustrated (as defined in [23]). Importantly, we demonstrate that minimally frustrated networks can be simplified—one can obtain multiple smaller networks that can recapitulate the behavior exhibited by the larger network. Since biological networks are minimally frustrated (shown in [23]), it is possible to develop network models that can recapitulate biological behavior without the need to incorporate all of the details of biological regulation.

The minimal frustration property described here is closely related to the near-monotone nature of biological networks characterized previously by Sontag and coworkers [41, 42]. Monotone systems are unlikely to exhibit chaotic behaviors and their response to perturbations is robust and predictable. The near-monotonicity of biological networks has been implicated in their dynamically stable behavior [42]. In predicting that the dynamics of minimally frustrated biological networks are largely one-dimensional, with steady states that exhibit biological gene expression patterns having very large basins of attraction, the present analysis provides far more functional insight than does the prediction of monotone dynamics. Interestingly, the mathematical framework for characterizing the monotonicity of dynamical systems provides a promising methodology for extending the concept of minimal frustration beyond biological regulatory networks to signaling and other biochemical networks. This idea will be explored in a future study.

As genome-wide transcriptional profiling became commonplace, first via microarray analysis and then via RNA-seq, it was recognized that gene expression datasets are effectively low-dimensional, as evidenced by the extensive covariation in the gene expression levels [43, 44]. The observed low-dimensionality has been attributed to the co-regulation of genes within regulatory modules, with the number of modules in the underlying network determining the dimensionality of the gene expression dataset [29]. Our analysis suggests that the regulatory network need not be modular to generate gene expression datasets that are low-dimensional: the random networks in our analysis can be more or less modular as compared to the biological network with similar topological features (Fig. S1). The steady-state expression profiles obtained from the biological networks are however always more one-dimensional as compared to random networks (Fig. 1). While selection for networks that exhibit low-dimensional steady-state expression patterns automatically results in the emergence of minimally frustrated networks (Fig. 2 a-b), such a selection pressure does not result in networks with higher modularity (Fig. S4). These results clearly establish that low-dimensional steady-state gene expression is a consequence of the minimal frustration property of the underlying regulatory network, independent of the network modularity.

Sethna and co-workers have previously shown that systems biology models are sloppy—the dynamical behavior of these models is dominated by a small number of combinations of kinetic parameters [45, 46]. This property makes dynamical models with poorly constrained kinetic parameters sufficient for recapitulating biological behavior and for making useful predictions. In the present study, we have shown that regulatory network models can recapitulate biological behavior and make useful predictions even if the network connection matrix *J* is poorly constrained: one need not have a complete and exact description of the regulatory network underlying a biological process to obtain a useful model of the process. Interestingly, while the sloppy parameter sensitivities reported by Sethna and co-workers are not limited to systems biology models and seem to extend to multi-parameter models in general [47, 48], the behavior reported in the present study is restricted to minimally frustrated networks.

Just as parameter fits even to comprehensive time series data fail to yield precise estimates of the underlying kinetic parameters due to the sloppy parameter sensitivities of systems biology models [46], our analysis suggests that collecting gene expression profiles at increasingly higher resolution and from more and more cells [49] is unlikely to yield more accurate biological regulatory networks. Pratapa *et al*. [50], benchmarking twelve different network inference algorithms on a variety of simulated and experimental gene expression datasets, found low stability in network prediction across datasets for the same biological process, and little agreement between the predictions by different algorithms for the same dataset. This is unsurprising in light of our observation that the expression profiles generated by biological networks and other minimally frustrated networks can be recapitulated by various simpler networks that lack several of the nodes and edges present in the original network (Fig. 3 and Fig. 4), as well as by lower resolution, coarse-grained networks that approximate the activity of several nodes by a single regulator (Fig. 5). Thus, the present study explains why the inference of gene regulatory networks from expression data remains a formidable challenge despite more than 20 years of research, and one that is unlikely to benefit from higher-resolution experimental data [50]. Instead of striving to obtain exact regulatory networks involved in establishing cell type-specific gene expression patterns by collecting higher-resolution data and employing advanced statistical techniques, efforts to understand cellfate choice must focus on building imperfect, predictive network models with rapidly verifiable predictions, and on carrying out experiments that are optimally designed to constrain the network connection matrix.

Our analysis shows that in the case of biological regulatory networks and in the case of minimally frustrated networks in general, steady-state expression patterns are largely preserved under progressive coarse graining of the network, a simplification procedure during which sets of network nodes are combined into single regulators (Fig. 5). Such behavior was previously reported for the small cell lung cancer (SCLC) network [16] analyzed in Fig. 1 [51]. This result provides a theoretical explanation for the popularity, despite the increasingly high resolution at which gene expression levels can be characterized by modern experimental techniques [49], of approaches involving the aggregation of genes into meaningful sets. One such approach is gene set enrichment analysis [52], whose popularity has endured the transition from microarray analysis to RNA-seq as the preferred method for transcriptome characterization. Previous studies have shown that gene pathways-based metrics that coarse grain the information contained in the expression levels of the constituent genes retain crucial information about the biological sample [31]. Note that the strategy to coarse grain a regulatory network described here is fairly simplistic, and proposed only as a sample strategy to demonstrate that the network behavior can be preserved under such a procedure. There exist numerous possibilities for vastly improving upon the present strategy; for example, by introducing an objective function that a network coarse-graining procedure must optimize. Strategies for coarse graining other biological network models including signaling networks and metabolic networks have been described elsewhere [53, 54] and could motivate improvements to the strategy introduced here for regulatory networks.

Finally, while previous studies have attributed the robustness of biological networks to the different topological features of these networks, our analysis posits a fairly simple explanation: the structure of biological networks is far more complex and requires far more information to describe as compared to the nature of the underlying biological process. Consequently, loss of features such as nodes and edges from the biological network description is unlikely to significantly affect the biological behavior. Since simple steady-state dynamics can emerge from a complex regulatory network only if the network is minimally frustrated, we posit that the minimal frustration property of biological networks is responsible for their functional robustness.

This work was supported by the National Science Foundation grant PHY-2019745.

## Appendix: Methods

### 1. Generation of random networks

Given a specific biological network, random networks were generated in a manner that preserved the number of nodes and edges in the network, the number of activating and inhibitory edges in the network, and the in-degree and out-degree of each network node. Starting with the biological network, we chose two edges in the network at random and switched their target nodes. This operation was repeated multiple times to obtain a random network with similar topological features as the given biological network. The behavior of many such random networks generated by starting from a given biological network has been analyzed in Fig. 1, Fig. S1, and Fig. S3.

### 2. Random circuit perturbation (RACIPE)

The random circuit perturbation (RACIPE) approach [27] was developed to analyze the robust dynamical features of a regulatory network when the exact kinetic parameters governing the dynamics are unavailable. Given a network connection matrix *J*, this approach generates an ensemble of kinetic parameter sets for the system of ODEs in Eq. 2. RACIPE generates the ensemble in a systematic fashion so that the dynamical network behavior over the ensemble can capture the full range of biological possibilities (see [27] for details). Each parameter set in the ensemble is comprised of 2*N* + 3*E* individual parameters, where *N* is the number of nodes in the regulatory network and E is the number of edges in the network (equal to the number of non-zero entries in the matrix *J*). For each parameter set in the ensemble, the ODE system in Eq. 2 is integrated numerically starting from multiple random initial conditions. The steady states obtained for a given parameter set in the ensemble by integration over a long time and for multiple initial conditions can then be analyzed to determine if the network behavior for the given parameter set is mono-stable, bi-stable, or multi-stable.

In the present study, to analyze the behavior of a regulatory network using the RACIPE approach, we generated an ensemble with 100 parameter sets. For each parameter set in the ensemble, we analyzed the dynamics starting from 100 randomly generated initial conditions (unless specified otherwise). The set of steady states thus obtained for a given network connection matrix *J* was analyzed in this study (using principal component analysis (PCA), hierarchical clustering, etc.). Note that we consider all the unique steady states obtained for each parameter set. Thus, say we consider an ensemble consisting of 10 parameter sets. Simulating the network dynamics for each parameter set starting from random initial conditions, we find that 5 of the parameter sets result in one unique steady state each, 2 result in two steady states each, and 3 result in three steady states each. Thus, the set of steady states analyzed will consist of a total of (5 × 1) + (2 × 2) + (3 × 3) = 18 states. Before analysis, the expression / activity level of each node in the set of steady states was log_2_ transformed.

All RACIPE simulations were carried out using the code available from https://github.com/simonhb1990/RACIPE-1.0.

### 3. Hierarchical clustering and heatmaps

We used the MATLAB*® [55] function `clustergram` [56] to analyze the underlying structure in the set of steady states obtained for different regulatory networks. The *M* × *N* matrix of the set of steady states obtained for a given network (as described in Appendix Sec. 2; M steady states for a network with N nodes) was used as the input to the `clustergram` function. This function returns a heatmap showing the expression levels of different nodes in the various steady states. Here, the expression level is indicated by the color (see color bar for each heatmap). In the heatmap returned by the `clustergram` function, steady states with similar expression levels of the different nodes are shown together. At the same time, nodes with similar expression levels across the different steady states are also shown together. We used the Euclidean distance between vectors as the distance metric for hierarchical clustering. In each figure showing one or more heatmaps, we note in the caption if both rows and columns of the steady-state matrix were hierarchically clustered or if clustering was only carried out along a single dimension.

### 4. Evolution simulations

For the evolution simulations in Fig. 2, we first generated a population of random networks. For this, we used the 26 node, 100 edge epithelial-mesenchymal (EMT) network [22] as the starting point and followed the procedure described in Appendix Sec. 1. Each network in the population of random networks thus generated had the same topological features as the EMT network. For the simulation in Fig. 2 a-b, we started with a population of 127 random networks (chosen for faster calculations on the computing cluster). For the simulation in Fig. 2 c-f, we started with a population of 500 random networks. At each time point (generation), we calculated two scores for each network in the population:

1. *Frustration score* — We simulated the network dynamics within a Boolean modeling framework (see [23] for details) starting from 50 random initial conditions. For each random initial condition, we simulated the network dynamics for 500 discrete time steps and calculated the frustration of the end state. Thus, we obtained 50 frustration values for each network. The minimum of these values was assigned as the frustration score of the network. A low value of the frustration score for a network will thus indicate that the network is more likely to be minimally frustrated.
2. *PC1 score* — We used RACIPE to determine the set of steady states a network can exhibit within our ODE-based modeling framework (see Appendix Sec. 2). We then carried out principal component analysis on the set of steady states obtained and determined the principal component that accounts for the greatest fraction of the variance in the steady states (i.e., the first principal component, or PC1). The variance accounted for by this principal component was defined as the PC1 score of the network. A high PC1 score for a network will thus imply that the network steady-state dynamics are simple and largely one-dimensional.

Depending on the selection pressure regime chosen, we defined the top 5% of networks in a generation as the 5% of networks with either the highest PC1 scores (when selecting for networks with simple, largely one-dimensional steady-state dynamics; Fig. 2 a-b) or the lowest frustration scores (when selecting for networks with minimal frustration; Fig. 2 c-f). These top 5% networks were then used to populate the next generation: we randomly chose one of the top 5% networks and mutated it with a probability 0.05; this operation (random choice followed by mutation with a certain probability) was repeated enough times to recover the original population size. The mutation step involved randomly choosing a couple of network edges and switching their targets. While carrying out the simulations under no selection pressure (black trajectory in Fig. 2 a-d), the entire population in a generation (instead of the top 5% networks) was used used to populate the next generation.

Fig. 2 a, c show the average PC1 score of the population at different time points (generations). Fig. 2 b, d show the average frustration score of the population at different time points (generations).

### 5. Node and edge deletion

While generating simpler networks via node or edge deletion, we defined the node and edge sensitivities as follows:

1. *Node sensitivity* — Simulating the network dynamics using RACIPE for 25 different parameter sets and 5 random initial conditions for each parameter set, the sensitivity for the node *j* was defined as:

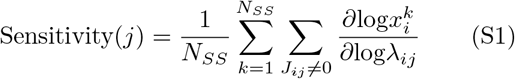
2. *Edge sensitivity* — Once again, simulating the network dynamics using RACIPE for 25 different parameter sets and 5 random initial conditions for each parameter set, the sensitivity for the edge *i* ← *j* was defined as:

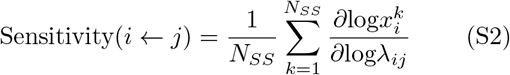

Here, *N_SS_* is the number of steady states obtained, *x^k^* is the expression / activity level of *i*^th^ node in the *k*^th^ steady-state, and λ_*ij*_ is a network kinetic parameter associated with the network edge *i* ← *j* (see Eq. 2).

In Fig. 3 and Fig. S5, the plot line labeled “Remove nodes / edges in order of sensitivity” was generated by deleting the node or edge with the lowest sensitivity identified at each step. The other plot lines in these figures were generated by deleting a randomly chosen node or edge at each step.

### 6. Network coarse graining

To simplify a given network via coarse graining, we used RACIPE (see Appendix Sec. 2) to obtain the set of steady states that can be exhibited by the network (100 parameter sets, 100 random initial conditions for each parameter set). We then used the MATLAB*® [55] function linkage [57] to carry out agglomerative hierarchical clustering of the network nodes, using the Pearson pairwise correlation calculated across the steady states as the distance metric: distance between nodes *i* and *j* was defined as 1 – corr(*i, j*). The function linkage returns a hierarchy (i.e., a tree) representing the agglomerative clustering of the network nodes. Pairs of nodes grouped at the lowest level of the tree were combined together to form a single regulator as per the following rules:

1. If any of the nodes being combined into a single regulator has a self-edge, the new regulator formed by combining this node with another node will also have a self-edge.
2. If there is an edge between the nodes being combined into a single regulator, the new regulator formed by combining the nodes will have a self-edge.
3. If there is a edge between any of the nodes combined and another network node (one that is not being combined into the same regulator; an outside node), there will be an edge between the new regulator formed by combining the nodes and the outside node.

The abovementioned rules could result in multiple edges between pairs of nodes in the coarse-grained network as well as multiple self-edges for some nodes. If all edges between a pair (or all self-edges for a given node) have the same sign (activating or inhibitory), we replace the multiple edges by a single edge. If the edges have different signs, we replace the multiple edges by a single edge which carries the sign of the majority of the multiple edges. If there is no such majority, we replace the multiple edges with a single edge with a randomly chosen sign.

### 7. Network inference from gene expression data

To infer from gene expression data a regulatory network that can recapitulate the gene expression patterns seen in breast tumor samples characterized by Terunuma *et al*. [33] and describe the cell-fate choice between high MYC-activation and low MYC-activation states, we started with the gene expression profiles of the 61 tumor samples from [33] and a list of 396 MYC activity-associated genes from a separate study [58]. Next, we obtained the transcription factors that regulate any of the MYC activity-associated genes using the TRRUST database [59]. The expression levels of the MYC activity-associated genes and those of the transcription factors involved in their regulation were used as input to the GRNBoost2 algorithm [34]. The algorithm returned a list of ordered node pairs with possible regulatory relationships. GRNBoost2 assigns a score to each ordered pairwise regulatory relationship and we chose the 500 most important relationships thus scored to obtain a preliminary regulatory network. Since GRNBoost2 does not assign a sign (activating or inhibitory) to the regulatory relationships it reports, we used the sign of the pairwise correlation between node expression levels in the gene expression dataset to determine the nature of the regulatory relationship: the relationship is considered to be activating if the correlation between node pairs is positive and inhibitory otherwise. Finally, we removed from the network any nodes that do not regulate another network node and those that are not regulated by any other network node. The final regulatory network thus obtained consisted of 138 nodes and 451 edges.

To obtain the random networks analyzed in Fig. 6, we used randomly shuffled gene expression profiles as inputs to the network inference procedure described above. Unlike the random networks obtained using the procedure described in Appendix Sec. 1, the random networks obtained here do not necessarily have the same number of nodes and edges, or the same topological features as the inferred MYC-activation network.

### 8. Data availability

The networks analyzed in the present study and those obtained during the various network simplification procedures, i.e., via node deletion, edge deletion, or coarse graining, are available online at https://github.com/st35/frustration-ODEs-modeling/tree/main/networkfilesas.topo files. In each tab-separated .topo file, the first column indicates the source node, the second column indicates the target node, and the third column indicates the edge type (1 for an activating edge, 2 for an inhibitory edge). In the .topo files for coarse-grained networks, the names of regulators combined into a single regulator are separated by two consecutive colons (::).

## Supplementary Figures

**FIG. S1.**
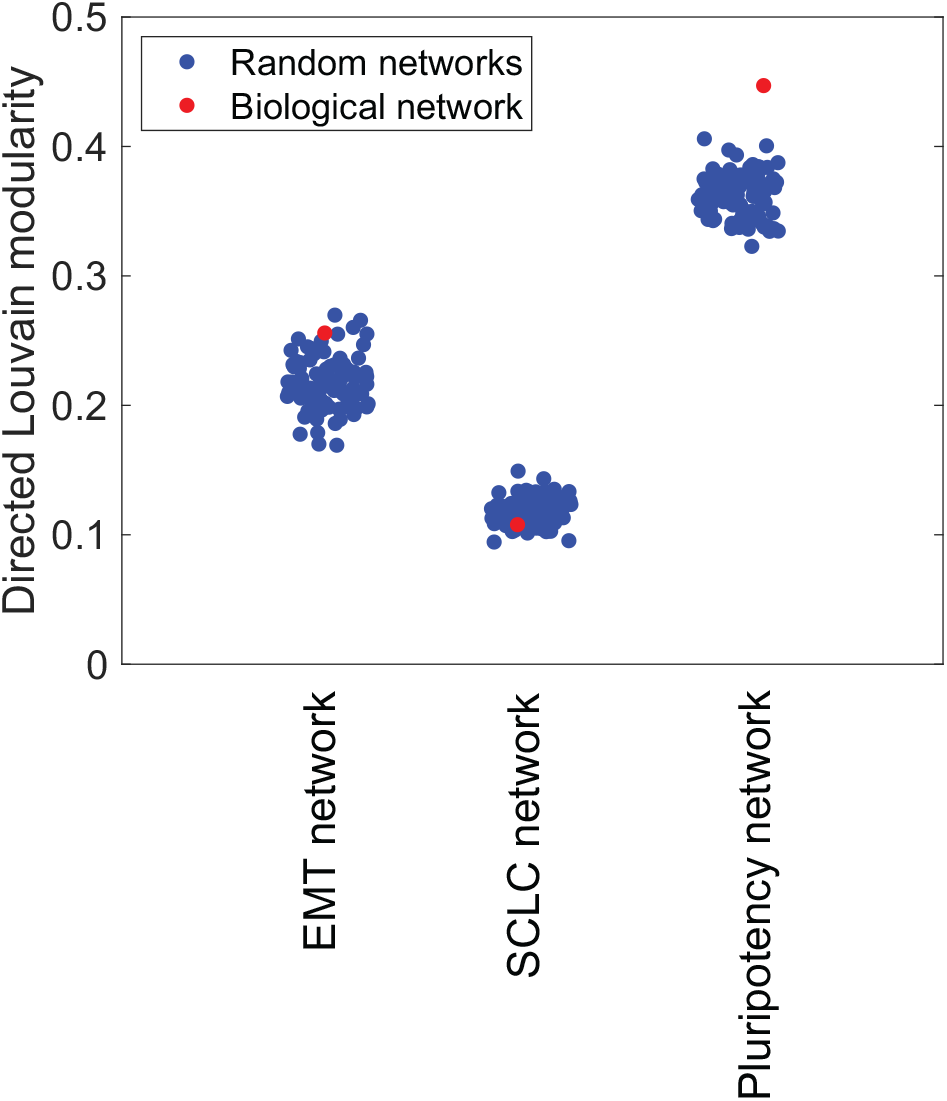
Directed Louvain modularity [26] of the different biological networks taken from the literature (red dots) and of random networks (blue dots) with similar topological features (generated using the approach in Appendix Sec. 1). Clearly, the random networks generated with either the epithelial-mesenchymal (EMT) network [22] or the small cell lung cancer (SCLC) network [16] as the starting point can be more or less modular as compared to the corresponding biological network. The pluripotency network is more modular than the random networks generated. Modularity values shown here were calculated using the code available at https://github.com/nicolasdugue/DirectedLouvain.

**FIG. S2.**
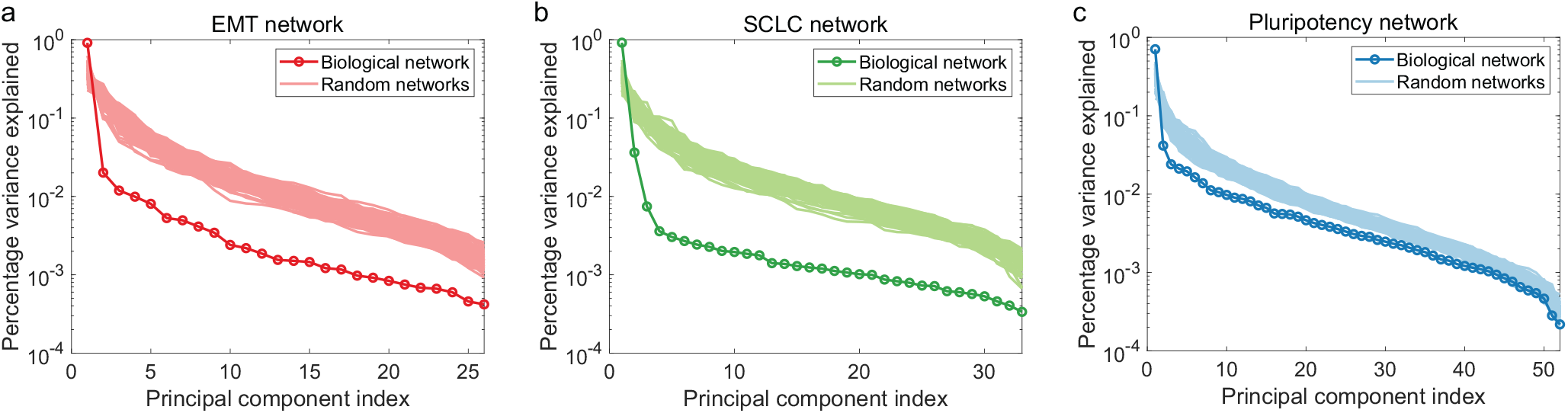
Percentage of variance in the steady states explained by different principal components for the epithelial-mesenchymal transition (EMT) network [22] (**a**), the small cell lung cancer (SCLC) network [16] (**b**), and the pluripotency network [15] (**c**). Note that the principal components were sorted in decreasing order of the percentage variance explained. Thus, in each case, the first principal component is the one that explains the greatest percentage of the variance in steady states. The sets of steady states for biological and random networks were obtained using the approach described in Appendix Sec. 2.

**FIG. S3.**
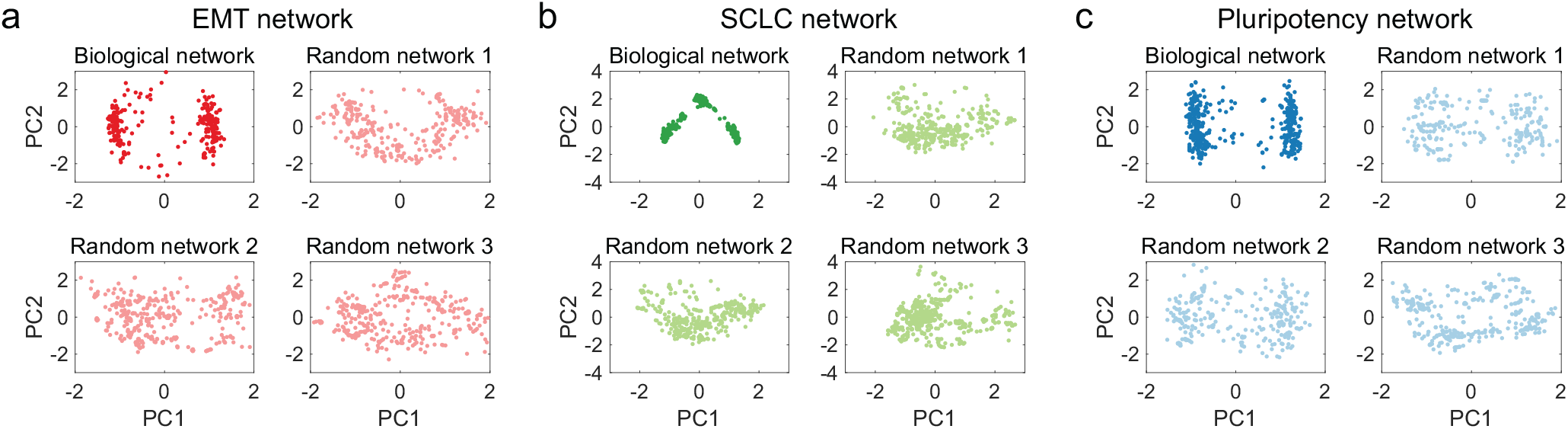
Steady states of biological and random networks shown in the space of the first two principal components (PC1 and PC2: the two components that account for the largest fraction of the variance in the steady states). In each plot, a dot corresponds to a single steady state. The top-left plot in each panel shows the behavior for a biological network (the epithelial-mesenchymal (EMT) network [22] in **a**, the small cell lung cancer (SCLC) network [16] in **b**, and the pluripotency network [15] in **c**). The remaining three plots in each panel show the behavior of random networks with the same topological features as the corresponding biological network.

**FIG. S4.**
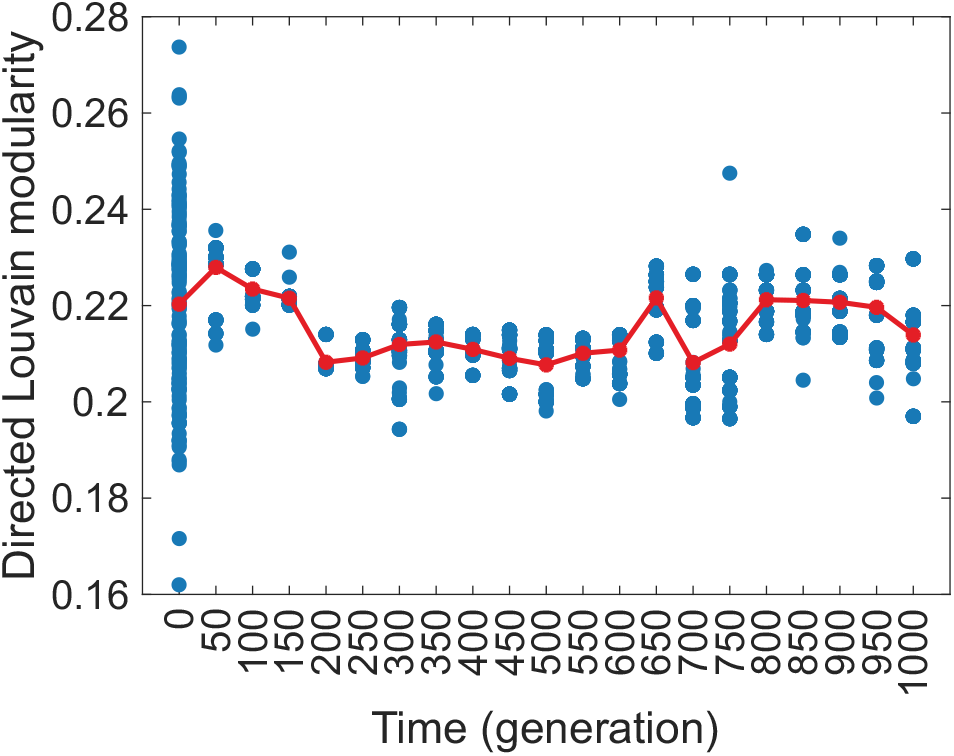
Directed Louvain modularity [26] of the networks in the population at different time points (generations), shown for the evolution simulation in Fig. 2 a-b, *i.e.*, under selection for networks with largely one-dimensional steady-state dynamics. Each blue dot shows the modularity of a single network in the population at a given time point. The red dots (and curve) show the modularity averaged over the networks in the population at a given time point. Clearly, while selecting for networks for which most of the steady-state variance can be explained by a single principal component, there is no automatic selection for networks with higher (or lower) modularity. Modularity values shown here were calculated using the code available at https://github.com/nicolasdugue/DirectedLouvain.

**FIG. S5.**
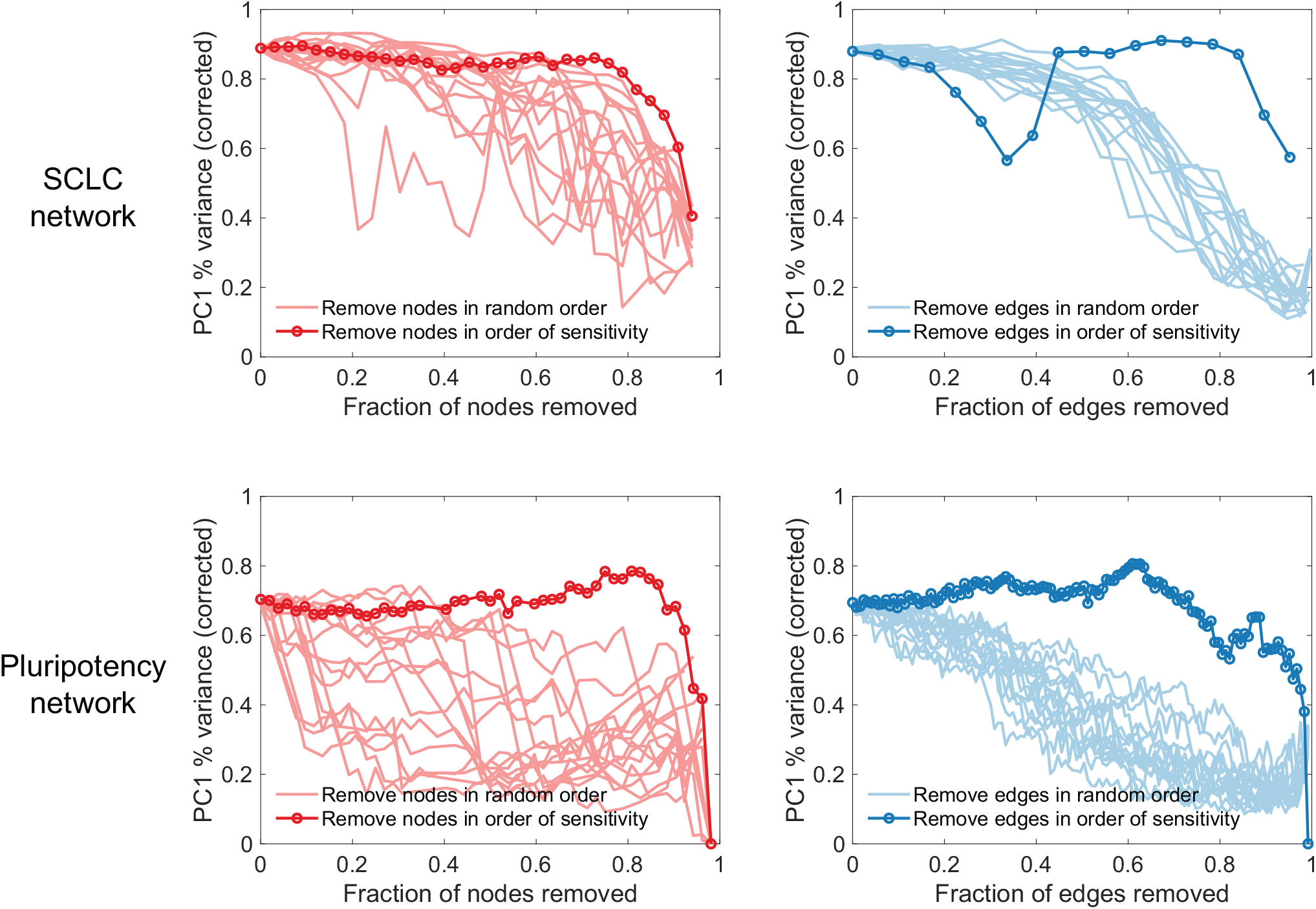
Same analysis as the one in Fig. 3 a-d, shown here for the small cell lung cancer (SCLC) network [16] (top row) and for the pluripotency network [15] (bottom row).

**FIG. S6.**
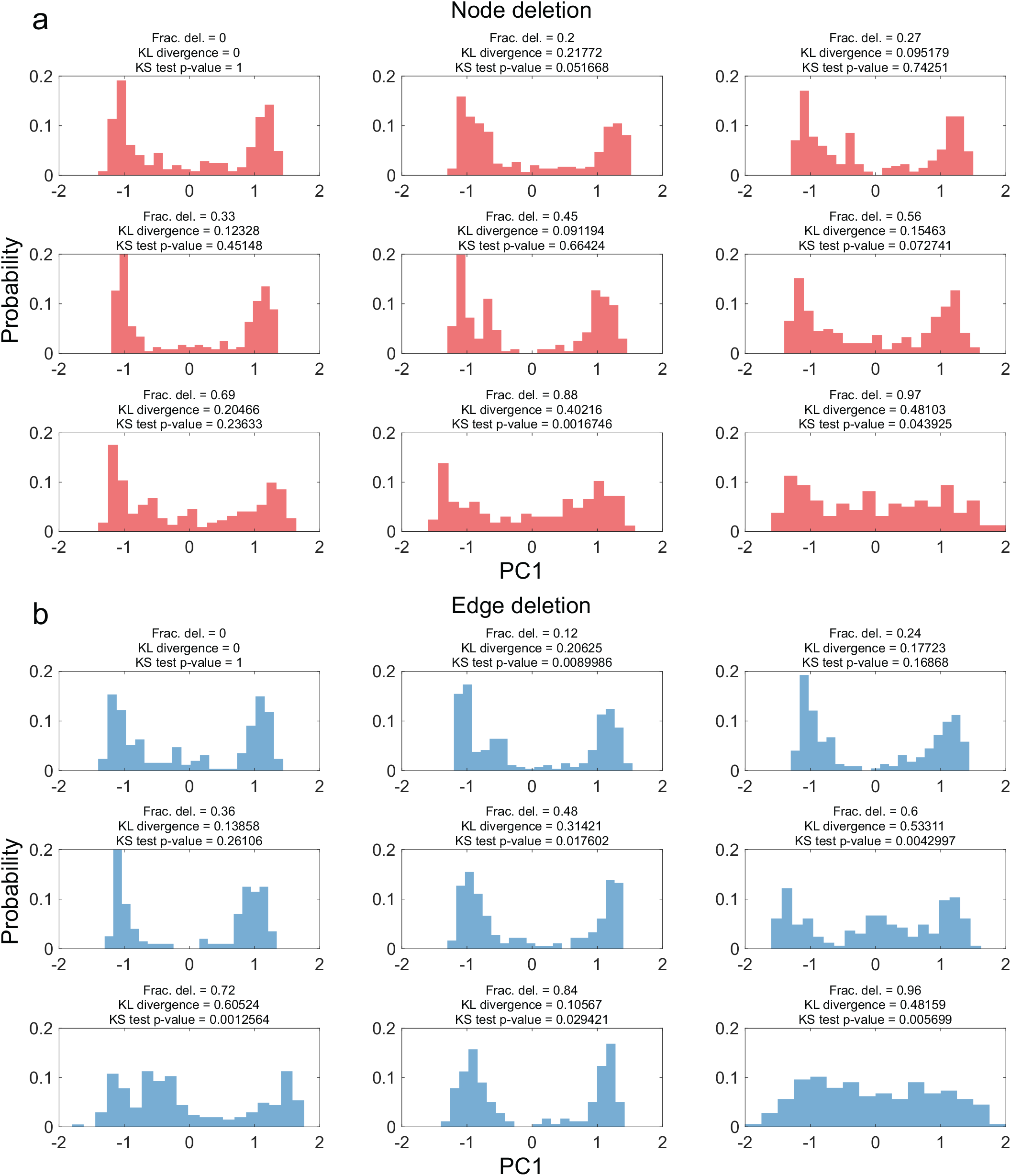
Same as Fig. 3 c-d, showing quantitative comparison between the first principal component (PC1) distribution obtained in the case of the 26 node, 100 edge minimally frustrated network (top-left plot in **a** and **b**) and the distribution for networks obtained by deleting different fractions of network nodes and edges (rest of the plots in **a** and **b**). KL divergence: Kullback-Leibler divergence; the KS (Kolmogorov-Smirnov) test p-value indicates the probability that the steady-state PC1 values for the original 26 node, 100 edge network and those for the simpler network (obtained by deleting a certain fraction of nodes or edges from the original network) are drawn from the same distribution.

**FIG. S7.**
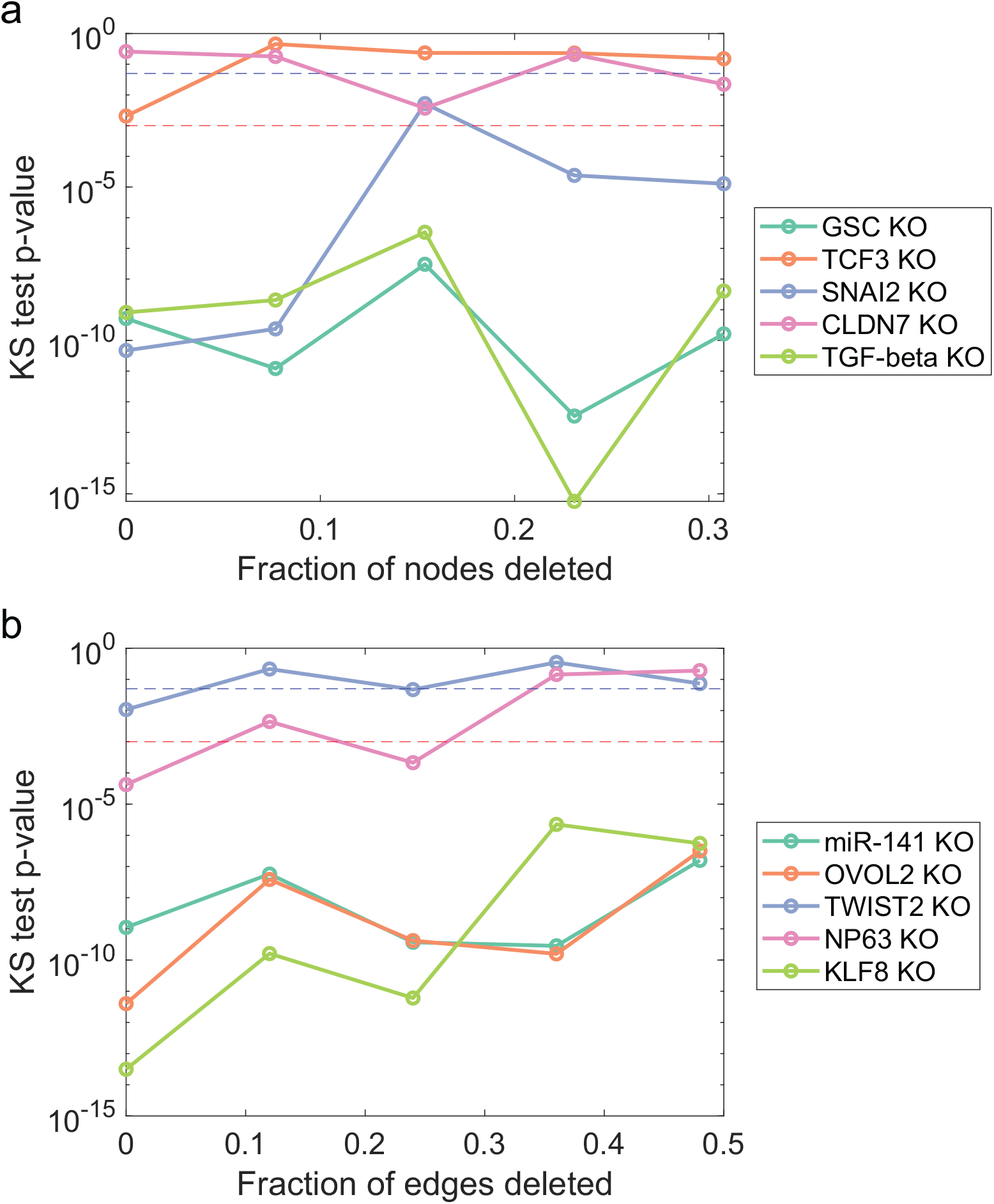
KS (Kolmogorov-Smirnov) test p-values for the comparison between the distribution of the first principal component (PC1) obtained after gene knockouts and the PC1 distribution in the control case (no gene knockout). We analyzed the gene knockout behavior for the 26 node, 100 edge EMT network [22] and for the simpler networks obtained via node deletion (panel **a**) or via edge deletion (panel **b**). These distributions are shown in Fig. 4 b (corresponding to panel **a** here) and Fig. 4 d (corresponding to panel **b** here). In both **a** and **b**, the blue dashed line indicates a p-value threshold of 0.05 while the red dashed line indicates a p-value threshold of 0.001. The KS test p-values here indicate the probabilities that the PC1 values in the control cases and those in the gene knockout cases are drawn from the same distribution.

**FIG. S8.**
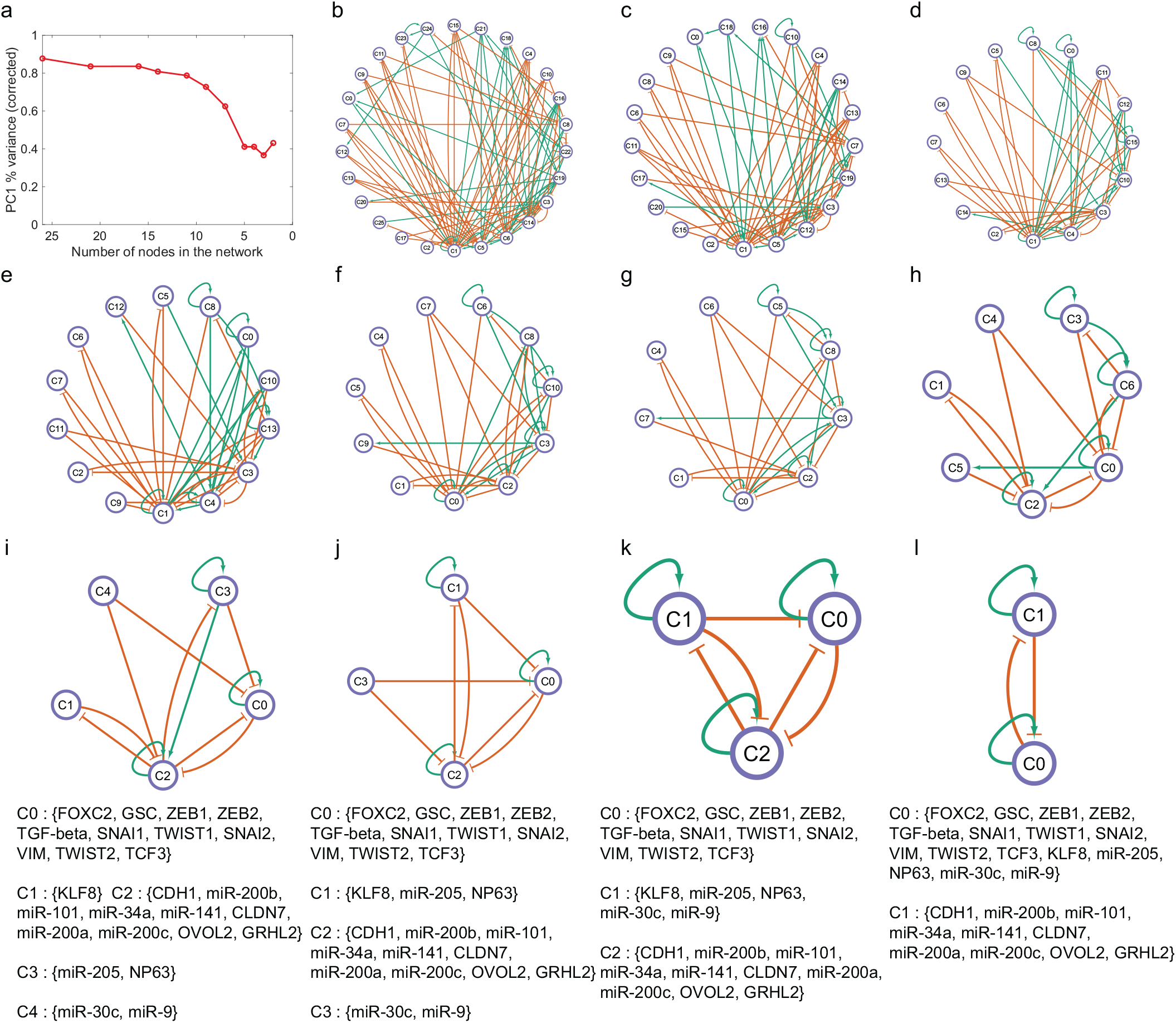
**a** Change in the percentage of the steady-state variance explained by the first principal component (PC1) as the 26 node, 100 edge epithelial-mesenchymal (EMT) network [22] is progressively coarse-grained. **b** The 26 node, 100 edge epithelial-mesenchymal (EMT) network [22]. **c**-**l** Networks obtained at different steps when the coarse-graining procedure (described in Appendix Sec. 6) is applied to the EMT network. For the networks in panels **i**-**l**, the nodes that have been clustered together into single regulators are also shown. Note that the coarse-graining procedure reduces the large and complex 26 node, 100 edge EMT network to a simple self-activating toggle switch between two sets of regulators: one consisting of well-known epithelial state factors (cluster C1 in panel **l**) and another consisting of well-known mesenchymal state factors (cluster C0 in panel **l**). This is consistent with previous analysis showing that a self-activating toggle switch can adequately model cell-fate choice between epithelial and mesenchymal states [60].

**FIG. S9.**
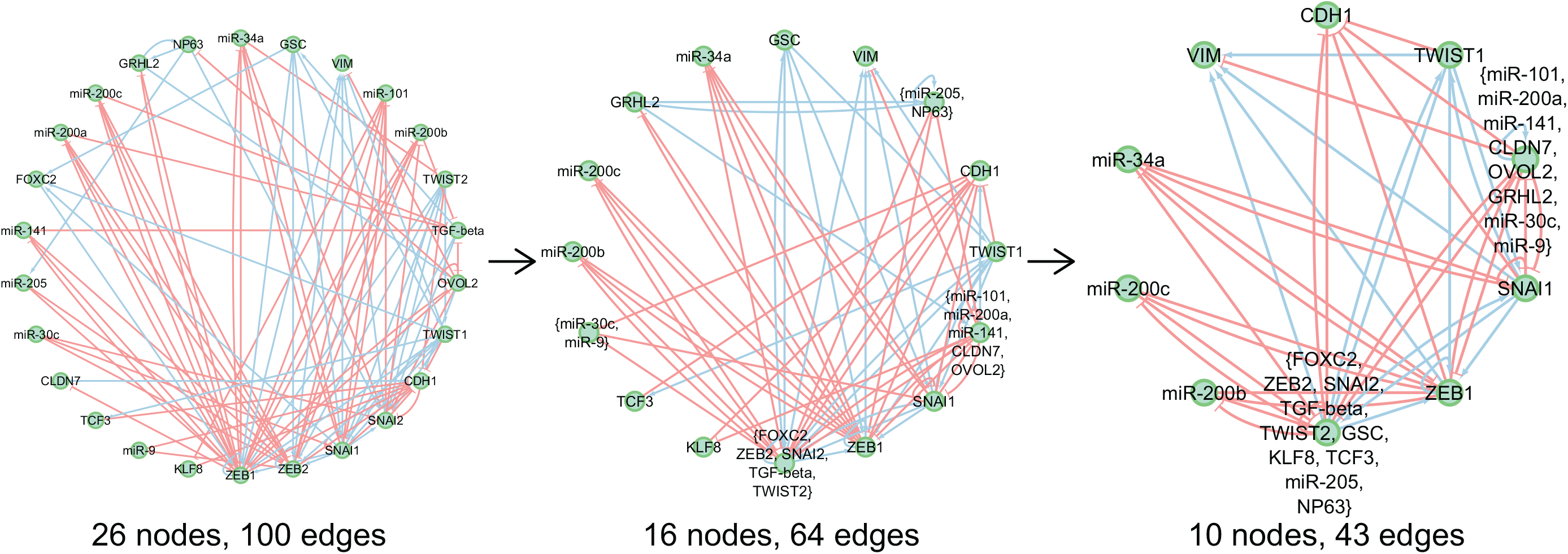
The original 26 node, 100 edge epithelial-mesenchymal (EMT) network [22] (left) and two of the simpler networks obtained via coarse graining (see Appendix Sec. 6 for the coarse-graining procedure).

**FIG. S10.**
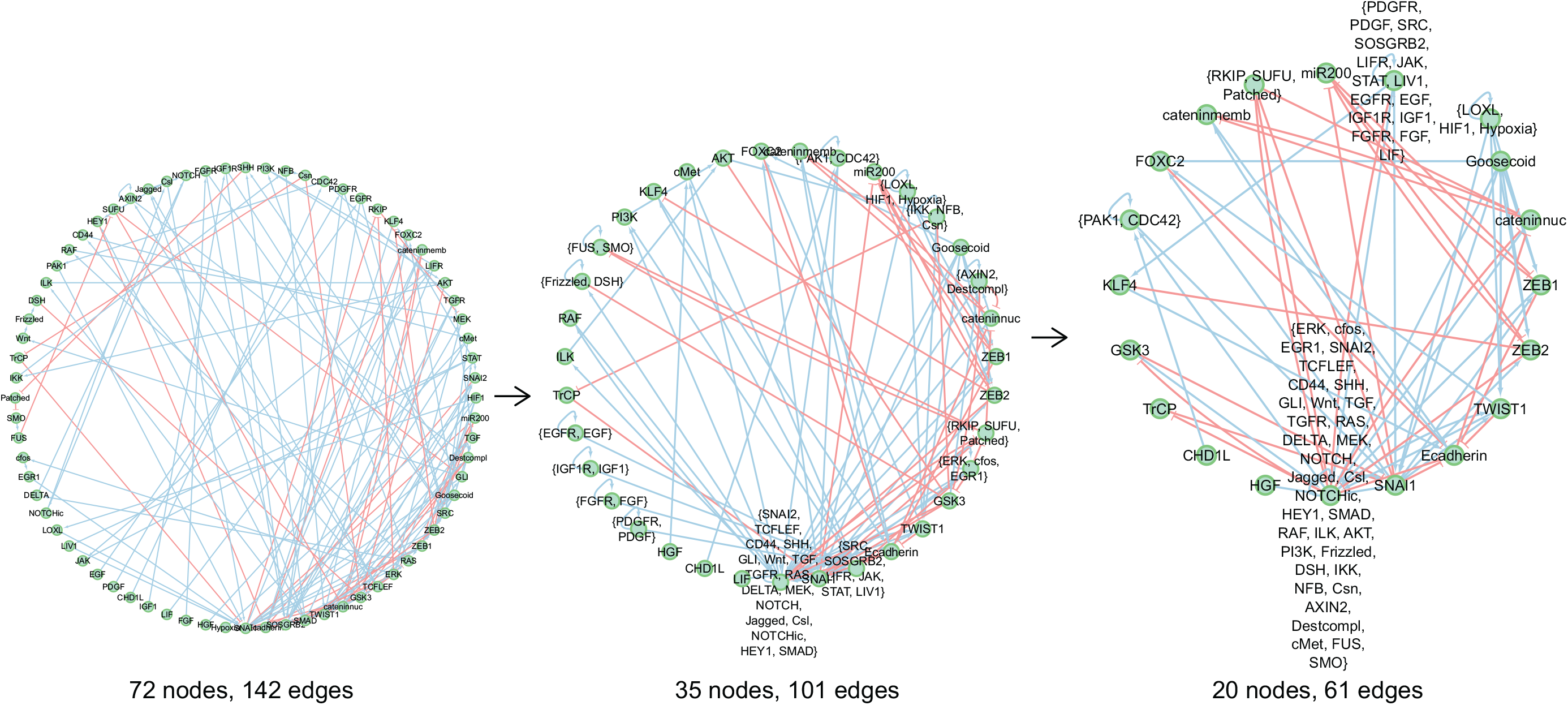
The original 72 node, 142 edge epithelial-mesenchymal (EMT) network [14] (left) and two of the simpler networks obtained via coarse graining (see Appendix Sec. 6 for the coarse-graining procedure).

**FIG. S11.**
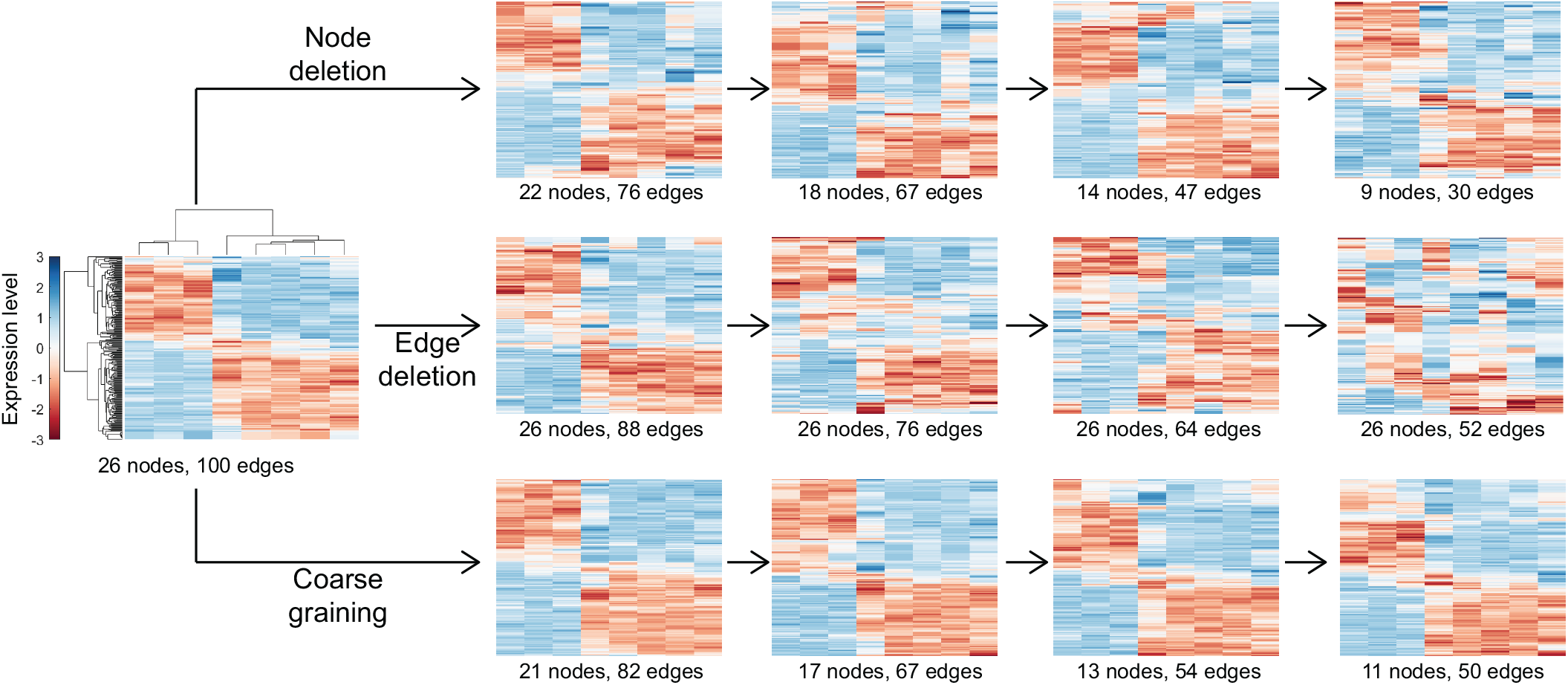
(Left) Steady-state expression patterns for the 26 node, 100 edge minimally frustrated network obtained via the evolution simulation in Fig. 2 **c**-**f** (selection for networks with minimal frustration). (Right) Simpler networks obtained via node deletion (top row), edge deletion (middle row), or coarse graining (bottom row) recapitulate the expression patterns exhibited by the larger evolved minimally frustrated network. Each heatmap shows the expression levels of the same eight nodes (randomly designated as nodes of interest) across the different steady states obtained for each network. Expression levels are indicated by the colors (see the color bar on the left). The simpler networks were obtained by deleting nodes and edges in random order while retaining the eight nodes of interest at each stage. While applying the coarse-graining procedure, the eight nodes of interest were not at any stage combined with any other network node. In all heatmaps, different steady states are shown along the rows while the network nodes are shown along the columns of the heatmaps. The heatmaps on the right were generated by hierarchically clustering the rows (*i.e.*, the steady states): each heatmap shows the network nodes (*i.e.*, the columns) in the same order as in the heatmap on the left (which was obtained by hierarchically clustering both the rows and the columns).

## Notes

### Competing Interest Statement

The authors have declared no competing interest.

## References

[1] T. Ideker and R. Nussinov, PLOS Comput. Biol. 13, 1 (2017).

[2] N. D. Price and I. Shmulevich, Curr. Opin. Biotechnol. 18, 365 (2007).

[3] R. Albert, The Plant Cell 19, 3327 (2007).

[4] T. Charitou, K. Bryan, and D. J. Lynn, Genet. Sel. Evol. 48, 27 (2016).

[5] F. J. Bruggeman and H. V. Westerhoff, Trends Microbiol. 15, 45 (2007).

[6] R. Albert, J. Cell Sci. 118, 4947 (2005).

[7] M. Aldana and P. Cluzel, Proc. Natl. Acad. Sci. U.S.A. 100, 8710 (2003).

[8] H. Yu and M. Gerstein, Proc. Natl. Acad. Sci. U.S.A. 103, 14724 (2006).

[9] R. Milo, S. Shen-Orr, S. Itzkovitz, N. Kashtan, D. Chklovskii, and U. Alon, Science 298, 824 (2002).

[10] J. E. Bailey, Nat. Biotechnol. 19, 503 (2001).

[11] R. Palmer, in Lectures in the Sciences of Complexity, edited by D. L. Stein (Addison-Wesley, Reading, MA, 1989) 1st ed., pp. 275–300.

[12] T. Enver, M. Pera, C. Peterson, and P. W. Andrews, Cell Stem Cell 4, 387 (2009).

[13] S. N. Steinway, J. G. Zañudo, W. Ding, C. B. Rountree, D. J. Feith, J. Loughran, Thomas P., and R. Albert, Cancer Res. 74, 5963 (2014).

[14] F. Font-Clos, S. Zapperi, and C. A. M. L. Porta, Proc. Natl. Acad. Sci. U.S.A. 115, 5902 (2018).

[15] R. Chang, R. Shoemaker, and W. Wang, PLOS Comput. Biol. 7, 1 (2011).

[16] A. R. Udyavar, D. J. Wooten, M. Hoeksema, M. Bansal, A. Califano, L. Estrada, S. Schnell, J. M. Irish, P. P. Massion, and V. Quaranta, Cancer Res, 77, 1063 (2017).

[17] S. Li, X. Zhu, B. Liu, G. Wang, and P. Ao, Oncotarget 6, 13607 (2015).

[18] O. Rios, S. Frias, A. Rodríguez, S. Kofman, H. Merchant, L. Torres, and L. Mendoza, Theor. Biol. Med. Model. 12, 26 (2015).

[19] O. Hobert, Science 319, 1785 (2008).

[20] M. Lu, M. K. Jolly, H. Levine, J. N. Onuchic, and E. Ben-Jacob, Proc. Natl. Acad. Sci. U.S.A. 110, 18144 (2013).

[21] X.-J. Tian, H. Zhang, and J. Xing, Biophys. J. 105, 1079 (2013).

[22] D. Jia, J. T. George, S. C. Tripathi, D. L. Kundnani, M. Lu, S. M. Hanash, J. N. Onuchic, M. K. Jolly, and H. Levine, Phys. Biol. 16, 025002 (2019).

[23] S. Tripathi, D. A. Kessler, and H. Levine, Phys. Rev. Lett. 125, 088101 (2020).

[24] R. Albert and J. Thakar, WIREs Syst. Biol. Med. 6, 353 (2014).

[25] P. Anderson, J. Less Common Met. 62, 291 (1978).

[26] N. Dugué and A. Perez, Directed Louvain: maximizing modularity in directed networks, Research Report (Universite d’Orléans, 2015).

[27] B. Huang, M. Lu, D. Jia, E. Ben-Jacob, H. Levine, and J. N. Onuchic, PLOS Comput. Biol. 13, 1 (2017).

[28] I. Pastushenko, A. Brisebarre, A. Sifrim, M. Fioramonti, T. Revenco, S. Boumahdi, A. Van Keymeulen, D. Brown, V. Moers, S. Lemaire, et al., Nature 556, 463 (2018).

[29] G. Heimberg, R. Bhatnagar, H. El-Samad, and M. Thomson, Cell Syst. 2, 239 (2016).

[30] J. Stelling, U. Sauer, Z. Szallasi, F. J. Doyle, and J. Doyle, Cell 118, 675 (2004).

[31] Y. Drier, M. Sheffer, and E. Domany, Proc. Natl. Acad. Sci. U.S.A. 110, 6388 (2013).

[32] A. P. Deshmukh, S. V. Vasaikar, K. Tomczak, S. Tri-pathi, P. den Hollander, E. Arslan, P. Chakraborty, R. Soundararajan, M. K. Jolly, K. Rai, et al., Proc. Natl. Acad. Sci. U.S.A. 118, e2102050118 (2021).

[33] A. Terunuma, N. Putluri, P. Mishra, E. A. Mathé, T. H. Dorsey, M. Yi, T. A. Wallace, H. J. Issaq, M. Zhou, J. K. Killian, et al., J. Clin. Investig. 124, 398 (2014).

[34] T. Moerman, S. Aibar Santos, C. Bravo González-Blas, J. Simm, Y. Moreau, J. Aerts, and S. Aerts, Bioinformatics 35, 2159 (2018).

[35] W. Saelens, R. Cannoodt, H. Todorov, and Y. Saeys, Nat. Biotechnol. 37, 547 (2019).

[36] K. R. Moon, J. S. Stanley, D. Burkhardt, D. van Dijk, G. Wolf, and S. Krishnaswamy, Curr. Opin. Syst. Biol. 7, 36 (2018).

[37] J. T. George, M. K. Jolly, S. Xu, J. A. Somarelli, and H. Levine, Cancer Res. 77, 6415 (2017).

[38] T. Z. Tan, Q. H. Miow, Y. Miki, T. Noda, S. Mori, R. Y.-J. Huang, and J. P. Thiery, EMBO Mol. Med. 6, 1279 (2014).

[39] L. A. Byers, L. Diao, J. Wang, P. Saintigny, L. Girard, M. Peyton, L. Shen, Y. Fan, U. Giri, P. K. Tumula, et al., Clin. Cancer Res. 19, 279 (2013).

[40] S. W. Ng, A. Mitchell, J. A. Kennedy, W. C. Chen, J. McLeod, N. Ibrahimova, A. Arruda, A. Popescu, V. Gupta, A. D. Schimmer, et al., Nature 540, 433 (2016).

[41] E. D. Sontag, Syst. Synth. Biol. 1, 59 (2007).

[42] A. Ma’ayan, A. Lipshtat, R. Iyengar, and E. Sontag, IET Syst. Biol. 2, 103 (2008).

[43] M. B. Eisen, P. T. Spellman, P. O. Brown, and D. Botstein, Proc. Natl. Acad. Sci. U.S.A. 95, 14863 (1998).

[44] S. Bergmann, J. Ihmels, and N. Barkai, Phys. Rev. E 67, 031902 (2003).

[45] K. S. Brown, C. C. Hill, G. A. Calero, C. R. Myers, K. H. Lee, J. P. Sethna, and R. A. Cerione, Phys. Biol. 1, 184 (2004).

[46] R. N. Gutenkunst, J. J. Waterfall, F. P. Casey, K. S. Brown, C. R. Myers, and J. P. Sethna, PLOS Comput. Biol. 3, 1 (2007).

[47] K. S. Brown and J. P. Sethna, Phys. Rev. E 68, 021904 (2003).

[48] B. B. Machta, R. Chachra, M. K. Transtrum, and J. P. Sethna, Science 342, 604 (2013).

[49] V. Svensson, R. Vento-Tormo, and S. A. Teichmann, Nat. Protoc. 13, 599 (2018).

[50] A. Pratapa, A. P. Jalihal, J. N. Law, A. Bharadwaj, and T. Murali, Nat. Methods 17, 147 (2020).

[51] L. Chauhan, U. Ram, K. Hari, and M. K. Jolly, eLife 10, e64522 (2021).

[52] A. Subramanian, P. Tamayo, V. K. Mootha, S. Mukherjee, B. L. Ebert, M. A. Gillette, A. Paulovich, S. L. Pomeroy, T. R. Golub, E. S. Lander, et al., Proc. Natl. Acad. Sci. U.S.A. 102, 15545 (2005).

[53] M. K. Transtrum and P. Qiu, Phys. Rev. Lett. 113, 098701 (2014).

[54] H. M. Rasool and S. H. Khoshnaw, arXiv preprint arXiv:2109.06566 (2021).

[55] MATLAB, version 9.10.0.1669831 (R2021a) Update 2 (The MathWorks Inc., Natick, Massachusetts, 2021).

[56] Object containing hierarchical clustering analysis data - MATLAB, https://www.mathworks.com/help/bioinfo/ref/clustergram.html.

[57] Agglomerative hierarchical cluster tree - MATLAB linkage, https://www.mathworks.com/help/stats/linkage.html.

[58] S. Chandriani, E. Frengen, V. H. Cowling, S. A. Pendergrass, C. M. Perou, M. L. Whitfield, and M. D. Cole, PLOS ONE, 1 (2009).

[59] H. Han, J.-W. Cho, S. Lee, A. Yun, H. Kim, D. Bae, S. Yang, C. Y. Kim, M. Lee, E. Kim, et al., Nucleic Acids Res. 46, D380 (2017).

[60] M. K. Jolly, S. C. Tripathi, D. Jia, S. M. Mooney, M. Celiktas, S. M. Hanash, S. A. Mani, K. J. Pienta, E. Ben-Jacob, and H. Levine, Oncotarget 7, 27067 (2016).

